# Spatial control of the APC/C ensures the rapid degradation of Cyclin B1

**DOI:** 10.1101/2023.10.26.564157

**Authors:** Luca Cirillo, Rose Young, Sapthaswaran Veerapathiran, Annalisa Roberti, Molly Martin, Azzah Abubacar, Camilla Perosa, Catherine Coates, Reyhan Muhammad, Theodoros I. Roumeliotis, Jyoti S. Choudhary, Claudio Alfieri, Jonathon Pines

## Abstract

The proper control of mitosis depends on the ubiquitin-mediated degradation of the right mitotic regulator at the right time. This is under the control of the anaphase promoting complex/cyclosome (APC/C) ubiquitin ligase that is regulated by the Spindle Assembly Checkpoint (SAC). The Checkpoint prevents the APC/C from recognizing Cyclin B1, the essential anaphase and cytokinesis inhibitor, until all chromosomes are attached to the spindle. Once chromosomes are attached, Cyclin B1 is rapidly degraded to enable chromosome segregation and cytokinesis. We have a good understanding of how the SAC inhibits the APC/C, but relatively little is known about how the APC/C recognises Cyclin B1 as soon as the SAC is turned off. Here, by combining live cell imaging, *in vitro* reconstitution, biochemistry, and structural analysis by cryo-electron microscopy, we provide evidence that the rapid recognition of Cyclin B1 in metaphase requires spatial regulation of the APC/C. Using fluorescence cross correlation spectroscopy, we find that Cyclin B1 and the APC/C primarily interact at the mitotic apparatus. We further show that this is because Cyclin B1, like the APC/C, binds to nucleosomes, and identify an ‘arginine-anchor’ in the N-terminus as necessary and sufficient for binding to the nucleosome. Mutating the nucleosome binding motif on Cyclin B1 reduces its interaction with APC/C and delays its degradation, and cells with the mutant, non-nucleosome-binding Cyclin B1 become aneuploid, demonstrating the physiological relevance of our findings. Together, our data demonstrate that mitotic chromosomes constitute a platform to promote the efficient interaction between Cyclin B1 and APC/C and ensure the timely degradation of Cyclin B1 and genomic stability.

## INTRODUCTION

Protein degradation imposes unidirectionality on the cell cycle through the rapid degradation of key mitotic regulators (e.g. Cyclin B and Securin) in metaphase. This ensures that chromosome segregation and mitotic exit are rapid and effectively irreversible, but the corollary is that to maintain genome stability, degradation must be coordinated with chromosome attachment to the mitotic apparatus. To achieve this, the Spindle Assembly Checkpoint (SAC) monitors kinetochore attachment to microtubules and prevents the Anaphase Promoting Complex / Cyclosome (APC/C) ubiquitin ligase from recognising Cyclin B and Securin until all chromosomes have attached to the spindle (reviewed in Alfieri et al., 2017; Yamano, 2019). Once all the chromosomes have attached to the spindle, the APC/C rapidly recognises Cyclin B1 and Securin to drive sister chromatid separation and exit from mitosis. Although we know many of the components of the machinery and their molecular structures, we do not understand how they interact in living cells to generate rapid and responsive pathways.

The APC/C is activated at nuclear envelope breakdown by the phosphorylation of the APC/C subunits Apc1 and Apc3 and dephosphorylation of its co-activator, CDC20 (Fujimitsu and Yamano, 2021; Kraft et al., 2003; Labit et al., 2012; Steen et al., 2008; Zhang et al., 2016; reviewed in Yamano, 2019). These enable CDC20 to bind and activate the apo-APC/C, but it is then kept in check by the SAC. The SAC inhibits the APC/C by generating the Mitotic Checkpoint Complex (MCC, reviewed in Lara-Gonzalez et al., 2021; McAinsh and Kops, 2023), that binds tightly to the APC/C as a pseudo-substrate inhibitor (Alfieri et al., 2016; Izawa and Pines, 2015, 2011). When bound to the MCC, APC/C is unable to recognise and ubiquitinate Cyclin B1 and Securin (Thornton and Toczyski, 2003) although it is still able to recognise early targets such as Cyclin A2 and Nek2A (Alfieri et al., 2020; den Elzen and Pines, 2001; Di Fiore and Pines, 2010; Geley et al., 2001; Zhang et al., 2019). Once all chromosomes are properly attached to spindle microtubules the SAC is silenced and the APC/C can ubiquitylate Cyclin B1 and Securin, whose proteolysis by the 26S proteasome activates the Separase protease (Holland and Taylor, 2006; Stemmann et al., 2001; Yu et al., 2021) to cleave cohesin and allow sister chromatids to separate.

Strict temporal inhibition of the APC/C is necessary to prevent premature chromosome separation that may lead to the gain or loss of chromosomes by the daughter cells, but single-cell imaging has shown that Cyclin B1 rapidly begins to be degraded as soon as all the chromosomes are attached to the spindle (Jackman et al., 2020), which ensures the rapid exit from mitosis and could conceivably be required for the fidelity of chromosome segregation. But how the APC/C rapidly recognises its substrates when the SAC is silenced is not understood. One potential clue comes from our previous work showing that Cyclin B1-GFP disappears first from chromosomes and centrosomes compared with the cytoplasm in HeLa cells (Clute and Pines, 1999). Moreover, fractionation experiments on HeLa cells show that the APC/C may be more active on chromatin (Sivakumar et al., 2014). In Drosophila, Cyclin B1-GFP first disappeared from centrosomes and then from the spindle and the chromosomes (Huang and Raff, 1999). These observations indicate that the APC/C might be spatially regulated in mitosis, which could ensure that Cyclin B1 is removed first from specific locations to coordinate the events of anaphase and mitotic exit. Yet despite these indications, direct evidence for spatial regulation of the APC/C has been missing.

Here, we provide direct evidence for the spatial control of APC/C activity. Using live cell imaging and fluorescence cross correlation spectroscopy (FCCS) we have assayed the binding between Cyclin B1 and the APC/C and found that the pattern of their interaction correlates with that of Cyclin B1 degradation. Moreover, we have identified the molecular mechanism promoting this localised interaction: we found that the N-terminal of Cyclin B1 contains a nucleosome-binding motif that regulates Cyclin B1 chromatin localisation. Preventing Cyclin B1 from binding to chromatin delays its degradation and cells become aneuploid; timely degradation can be restored by recruiting a non-nucleosome-binding Cyclin B1 mutant back to the chromatin. Our data indicate that nucleosomes facilitate the interaction between Cyclin B1 and APC/C to promote the rapid degradation of Cyclin B1 during metaphase and safeguard genomic stability.

## RESULTS

### Cyclin B1 degradation is spatially regulated during metaphase

We used CRISPR/Cas9 gene editing to tag both alleles of Cyclin B1 in human retinal pigment epithelial (RPE-1) cells with the mEmerald fluorescent protein (Barbiero et al., 2022; Cubitt et al., 1998). As we and others previously reported, live-cell imaging revealed that Cyclin B1 localised to specific structures in mitotic cells, including the spindle microtubules, spindle caps, and chromosomes (Bancroft et al., 2020; Bentley et al., 2007; Huang and Raff, 1999; Jackman et al., 1995; Pines and Hunter, 1991), and Cyclin B1 fluorescence disappeared rapidly at the start of metaphase when the SAC is turned off (Clute and Pines, 1999; Hagting et al., 1998; Jackman et al., 2020, 2003). In agreement with our previous results using ectopically expressed Cyclin B1-FP (Clute and Pines, 1999), measuring the fluorescence intensity of Cyclin B1 at different subcellular locations showed that Cyclin B1 disappeared first from the chromosomes, spindle and centrosomes and only later in the cytoplasm (Fig. 1A, B).

**Figure 1.**
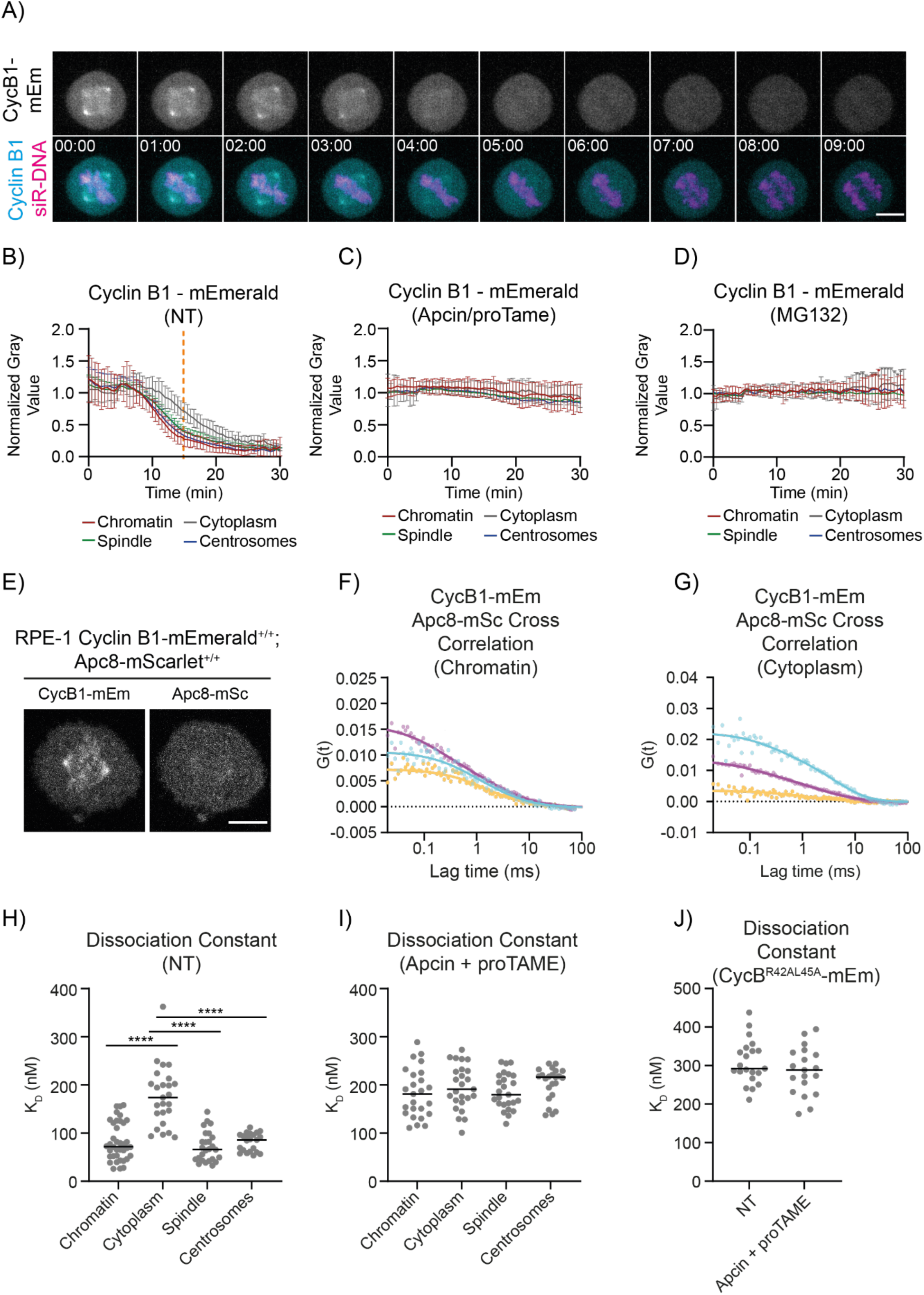
Cyclin B1 degradation is spatially regulated. A) Representative maximum projections of fluorescence confocal images over time of an RPE-1 CCNB1-mEmerald^+/+^ cell progressing through mitosis. Time is expressed as mm:ss. Scale bar correspond to 10 μm. B-D) Quantification of Cyclin B1 fluorescence levels (normalised Raw Integrated Density, RID) over time in RPE-1 CCNB1-mEmerald^+/+^ cells. n ≥ 18 cells per condition, N = 3 independent experiments. In this and all following graphs, mean ± Standard Deviation (SD), unless specified otherwise. E) Representative fluorescence confocal images of RPE-1 CCNB1-mEmerald^+/+^; APC8-mScarlet^+/+^ cells. Scale bar corresponds to 10 μm. F-G) Representative graph of the autocorrelation of Cyclin B1-mEmerald and APC8-mScarlet and the cross-correlation between the two at the Chromatin (F) and in the Cytoplasm (G). In this and all following FCCS graphs, dots = data, lines = fit; Cyan = Cyclin B1-mEmerald, magenta = APC8-mScarlet, yellow = Cross-Correlation between Cyclin B1-mEmerald and A8-mScarlet. H-J) Dot plots representing the K_D_ between endogenous Cyclin B1-mEmerald and APC8-mScarlet in the indicated conditions. n ≥ 21 cells per condition, N = 3 independent experiments. In this and all following K_D_ dot plots, each dot correspond to a single measurement, horizontal black lines represent median values. In this and all following figures, the dotted vertical line indicates anaphase, dotted horizontal line indicate 0, CycB1 = Cyclin B1, mEm = mEmerald, mSc = mScarlet, see table S1 for statistical tests and p-values.

Two non-exclusive mechanisms could account for the faster disappearance of Cyclin B1 in specific places: Cyclin B1 could be rapidly displaced, or it could be proteolysed *in situ*. To distinguish between these two possibilities, we measured the disappearance Cyclin B1 fluorescence in cells treated either with MG132 to inhibit the proteasome or with a combination of the APC/C inhibitors APCin and proTame (Sackton et al., 2014; Verma et al., 2004) to inhibit ubiquitylation. Both treatments stabilised Cyclin B1 levels and arrested cells in metaphase, as previously reported (Sackton et al., 2014; Zeng et al., 2010), and Cyclin B1 levels remained constant at all the subcellular locations tested (Fig. 1C, D). This indicated that the pattern of Cyclin B1 disappearance in metaphase is linked to its degradation and not to its rapid displacement.

### Analysing Cyclin B1-APC/C interaction by Fluorescence Cross Correlation Spectroscopy

Our results indicated that the proteolysis of endogenous Cyclin B1 is spatially regulated, therefore we set out to determine the mechanism responsible. This could originate from the differential activity of either the APC/C or the proteasome, or both. The APC/C was previously reported to be more active at centrosomes (Kraft et al., 2003) or chromosomes (Sivakumar et al., 2014; Topper et al., 2002), although these studies came to contradictory conclusions; therefore, we focused first on the interaction between Cyclin B1 and the APC/C. To measure protein-protein interactions with subcellular resolution in living cells, we turned to fluorescence cross correlation spectroscopy (FCCS), which we recently validated as a tool to study the cell cycle machinery (Barbiero et al., 2022). FCCS requires that the whole population of proteins under investigation be fluorescently tagged, therefore we used CRISPR/Cas9^D10A^ gene-editing (Shen et al., 2014) to introduce the mScarlet (Bindels et al., 2017) fluorescent protein into both alleles of the APC/C subunit APC8 in our RPE-1 CCNB1-mEmerald^+/+^ cells and the parental RPE-1 cells. PCR and sequencing confirmed biallelic insertion in three independent clones of both the parental RPE-1 cells and RPE-1 CCNB1-mEmerald^+/+^ cells (data not shown).

We first characterised the APC8-mScarlet cells to determine whether the tag perturbed APC/C function. APC8-mScarlet was enriched in the nucleus of interphase cells; during mitosis it was weakly enriched at centrosomes but otherwise homogeneous throughout the cell (Fig. 1E, S1A, B), in agreement with previous immunofluorescence data of other APC/C subunits (Acquaviva and Pines, 2006; Huang and Raff, 2002; Kraft et al., 2003; Tischer et al., 2022; Tugendreich et al., 1995). The mScarlet tag did not significantly alter mitotic timing in either unperturbed mitosis or in cells treated with low doses of the microtubule poison paclitaxel to prolong the SAC (Fig. S1C, Collin et al., 2013). Similarly, tagging APC8 had no significant effect on cell growth rate or ploidy (Fig. S1D). Although immunoblotting indicated that the protein levels of APC8-mScarlet were about 70% lower than wild type APC8 in parental cells (Fig. S1 E, F) this did not significantly affect the rate of Cyclin B1-mEmerald degradation in mitosis (Fig. S1H). To assess APC8 incorporation into the APC/C, we immunodepleted APC4 and measured how much APC8 was not co-immunoprecipitated and found there was no difference between APC8 and APC8-mScarlet (Fig. S1E, F). This indicated that the mScarlet tag did not interfere with APC8 incorporation into the APC/C. In agreement with this, FCS analysis of APC8-mScarlet in interphase and mitosis indicated that APC8-mScarlet behaved as a single, slow diffusing species with a diffusion coefficient of 6.435 ± 1.789 µm^2^ s^−1^ (Mean ± SD) and hydrodynamic radius of ∼ 14 nm which is comparable to the theoretical hydrodynamic radius of APC/C (Fig. S1G, Alfieri et al., 2016; Chang et al., 2015; Yamaguchi et al., 2016). We concluded that APC8 tagged with mScarlet was a valid reporter for the APC/C.

### D-box dependent Cyclin B1 – APC/C interaction is favoured in the spindle

Having validated cell lines where both alleles of Cyclin B1 and of APC8 were tagged with fluorescent proteins, we were able to use FCCS to study their interaction during cell division (Fig. 1E). In agreement with its SAC-dependent degradation there was no cross-correlation between Cyclin B1 and the APC/C in G2 phase cells but there was a strong interaction in metaphase cells (Fig. 1F, G, S2A). We used these data to calculate the dissociation constant (K_D_) between Cyclin B1-mEmerald and APC8-mScarlet at different subcellular locations and found that the interaction between Cyclin B1 and the APC/C had lower K_D_ at the chromatin, spindle and spindle poles than in the cytoplasm (Fig. 1F, G, H, S2B, C). We obtained similar results in cells arrested in metaphase with the proteasome inhibitor MG132 (Fig. S2D). To validate our FCCS assay we measured cross correlation in the presence of the small molecule inhibitors APCin and proTAME that impair CDC20 substrate recognition and CDC20 binding to the APC/C, respectively (Sackton et al. 2014). In the presence of APCin and proTAME, the K_D_ between Cyclin B1-mEmerald and APC8-mScarlet in the spindle, on chromosomes and centrosomes (Fig. 1I) increased to a level similar to that found in the cytoplasm of either treated or untreated cells (Fig. 1I).

The residual cross-correlation in the presence of APCin and proTAME indicated that there was a second, lower affinity mode of binding between Cyclin B1 and the APC/C, which was consistent with Cyclin B1-Cdk1-Cks binding to phospho-APC/C through the Cks protein (Kraft et al., 2003; Patra and Dunphy, 1996; Rudner and Murray, 2000; Shteinberg and Hershko, 1999; Sudakin et al., 1997; van Zon et al., 2010). In agreement with this, there was low level cross-correlation between Cyclin B1-mEmerald and APC8-mScarlet in prometaphase cells (Fig S2E, F), and this interaction was independent of the Cyclin B1 D-box (Fig. 1J, S2G) and unaffected by treatment with a combination of APCin and proTAME (Fig. 1J).

We concluded that FCCS revealed two major modes of binding between Cyclin B1 and APC/C in dividing RPE-1 cells: a high affinity, D-box-dependent interaction that was spatially constrained to the spindle and chromosomes, and temporally restricted to metaphase; plus a second, D box-independent, lower affinity interaction in both prometaphase and metaphase cells that was likely dependent on Cks.

### Chromatin is the major site of Cyclin B1 degradation in human cells

Our data indicated that Cyclin B1 degradation and its interaction with the APC/C was favoured on the chromosomes, spindle and centrosomes, but the relative contribution of each subcellular location to Cyclin B1 degradation was unclear. To answer this, we asked whether Cyclin B1 would disappear faster on chromosomes spatially separated from the metaphase plate. We generated cells in which some chromosomes remained at the spindle poles by inhibiting the CENP-E (Centromeric Protein E) kinesin motor protein that is required for these chromosomes to congress to the metaphase plate (Fig. 2A, Bennett et al., 2015). If individual chromosomes are important for Cyclin B1 degradation, then the degradation of Cyclin B1 should be faster on polar chromosomes than in the surrounding cytoplasm. Cells treated with a CENP-E inhibitor arrested in mitosis with polar chromosomes and stable levels of Cyclin B1 (Fig. 2B). When we treated CENP-E inhibited cells with the MPS1 inhibitor Reversine (Chen et al., 2004) to inactivate the SAC, Cyclin B1 disappeared earlier from polar chromosomes than it did from the surrounding cytoplasm (Fig. 2B). The decay of Cyclin B1 at polar chromatids was indistinguishable from that of aligned chromosomes.

**Figure 2.**
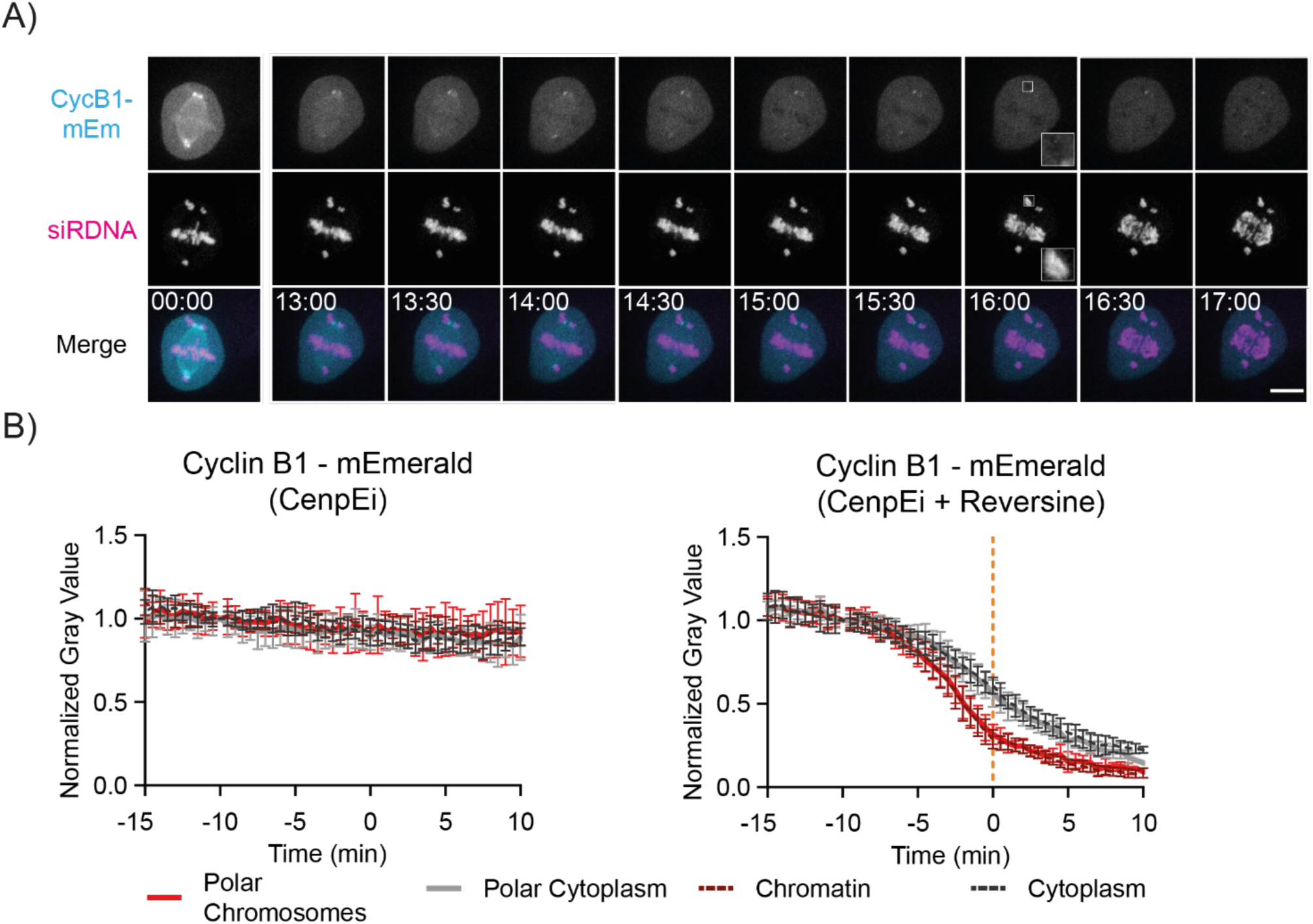
Chromatin is a major site for Cyclin B1 degradation in RPE-1 cells. A) Representative fluorescence confocal images over time of a RPE-1 CCNB1-mEmerald^+/+^ cell progressing through mitosis after a treatment with CenpEi and Reversine. Time is expressed as mm:ss. Scale bar corresponds to 10 μm. B) Quantification of normalised Cyclin B1 fluorescence levels over time in RPE-1 CCNB1-mEmerald^+/+^ cells after treatment with CenpEi inhibitor (left) or a combination of CenpEi and Reversine (Right). n = 11 cells per condition, N = 3 independent experiments.

This result demonstrated that the chromosomes themselves, rather than their relative position in the cell, facilitated Cyclin B1 degradation in metaphase and represent a major site of Cyclin B1 degradation.

### The APC3 loop is predicted to bind the nucleosome acidic patch

Our data supported the conclusion that chromosomes are the preferential site of Cyclin B1 interaction and degradation. A previous large-scale study found that the APC/C associates with nucleosomes via their acidic patch (Skrajna et al., 2020), which could be relevant to how chromosomes facilitate Cyclin B1 degradation. We tested whether APC/C could bind nucleosomes using an Electrophoretic Mobility Shift Assay with recombinantly reconstituted APC/C-CDC20 and Nucleosome Core Particles (NCP). (Fig. 3A), increasing concentrations of APC/C-CDC20 with an N-terminal fragment of Cyclin B1 (Cyclin B1^NTD^) shift the NCP, indicative of the formation of an APC/C-CDC20:NCP complex. To map the subunit of the APC/C responsible for the interaction with NCP, we used Cross-Linking mass spectrometry (XL-MS) with a complex of APC/C-CDC20-Cyclin B1^NTD^ and the NCP. In addition to the expected cross-links within the APC/C-CDC20 and within the NCP (Fig. S3 A, B), we identified two cross-links from the APC/C to the NCP (Fig. 3B): K390 from the APC3 loop region crosslinked to residues 96 and 106 of H2A and H2B respectively, which are proximal to the acidic patch of the nucleosome. Sequence alignment of the APC3 loop showed a conserved RxxRL motif (Fig. 3C), which is predicted to bind into the acidic patch by Alphafold multimer (Fig. 3D).

**Figure 3.**
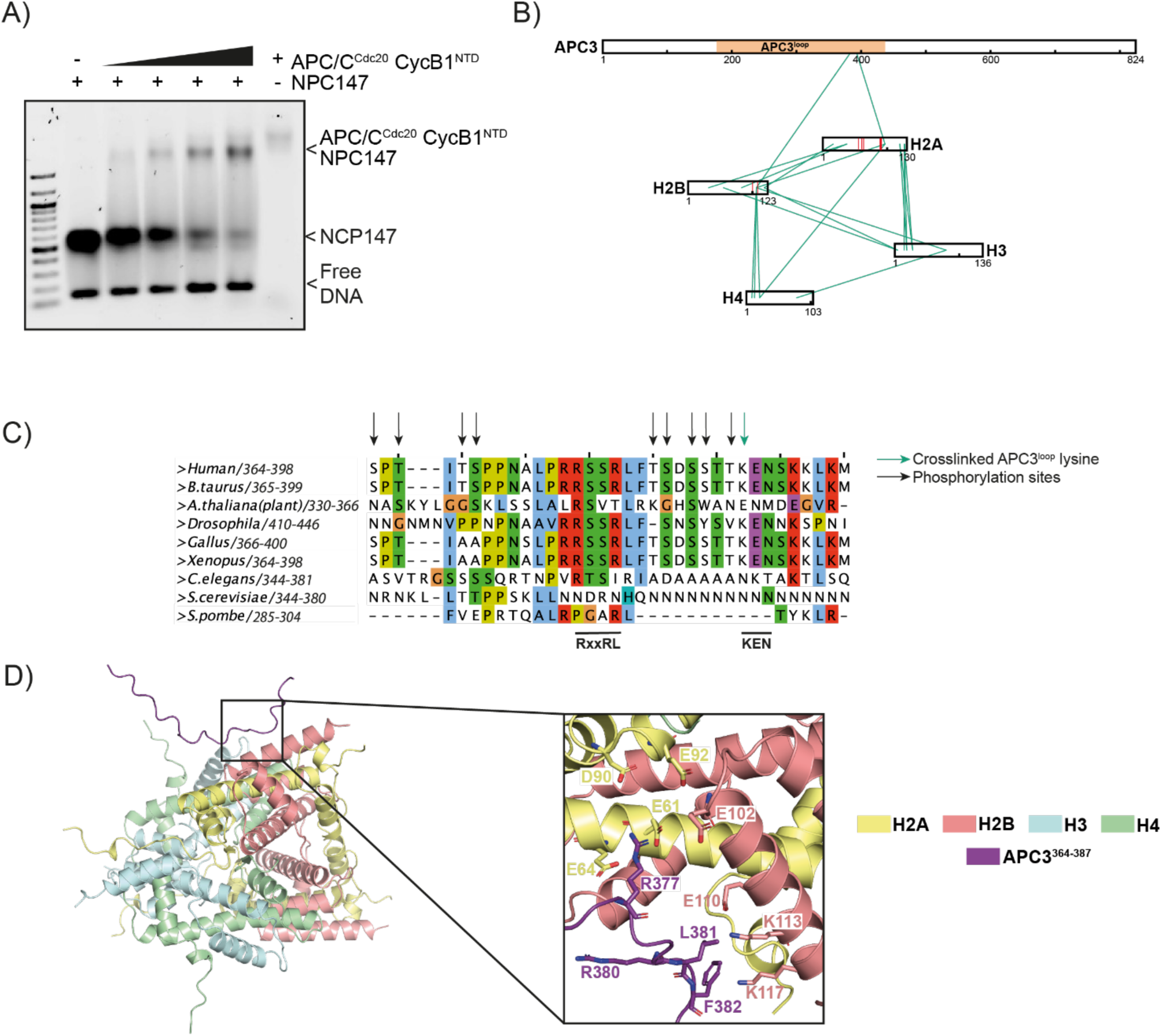
The APC3 loop of APC/C is predicted to interact with the nucleosome acidic patch. A) EMSA of the APC/C-CDC20-Cyclin B1^NTD^ and the NCP 147. N = 3 independent experiments. B) Representation of the heteromeric crosslinks found between the APC3 loop and the acidic patch of the NCP. C) A sequence alignment of the C-terminal end of human APC3 loop with homologous sequences in other species. APC3 K390 which crosslinks with the acidic patch is marked with a green arrow. Previously identified phosphorylation sites are marked with a black arrow. D) Alphafold multimer prediction of the interaction between the APC3 loop(364-387) and the histone octamer.

### N-terminus of Cyclin B1 directly interacts with the nucleosome acidic patch

Having established how the APC/C is likely to bind nucleosome, we investigated the interaction between Cyclin B1 and chromosomes. We focused on the N-terminus of Cyclin B1 because we had shown that the N-terminal 9 amino acids of Cyclin B1 were required for rapid degradation (Matsusaka et al., 2014), and work from the King lab demonstrated that arginines in the N-terminus of Cyclin B1 were important for chromosome recruitment (Bentley et al., 2007; Pfaff and King, 2013).

The sequence of the first seven amino acids of Cyclin B1 are conserved and resemble an ‘arginine anchor’ motif (Fig. 4A, B) that is used by several proteins to bind to the nucleosome acidic patch (reviewed in Kalashnikova et al., 2013; Paul, 2021). We used an EMSA to test the possibility that Cyclin B1 could bind directly to nucleosomes. (Fig. 4C, S3C). Adding increasing amounts of Cyclin B1^WT^ to the nucleosomes shifted the DNA-nucleosome band towards higher molecular masses, indicating the formation of a complex (Fig. 4C, S3C). We obtained similar results using an N-terminal peptide containing the first 23 AA of Cyclin B1 (Fig. S3D). We tested whether this interaction was mediated by the putative arginine anchor in the amino terminus of Cyclin B1 by mutating the arginine residues and observed a significant decrease in the binding affinity, in charge-substitution mutants (Cyclin B1^4E7E^, Fig. 4C, S3D), or when the first 9 AA of Cyclin B1 were deleted (CyclinB1^Δ9^) or amino acids 3 to 7 were mutated to alanine (Cyclin B1^3A5^, Fig. S3E, F).

**Figure 4.**
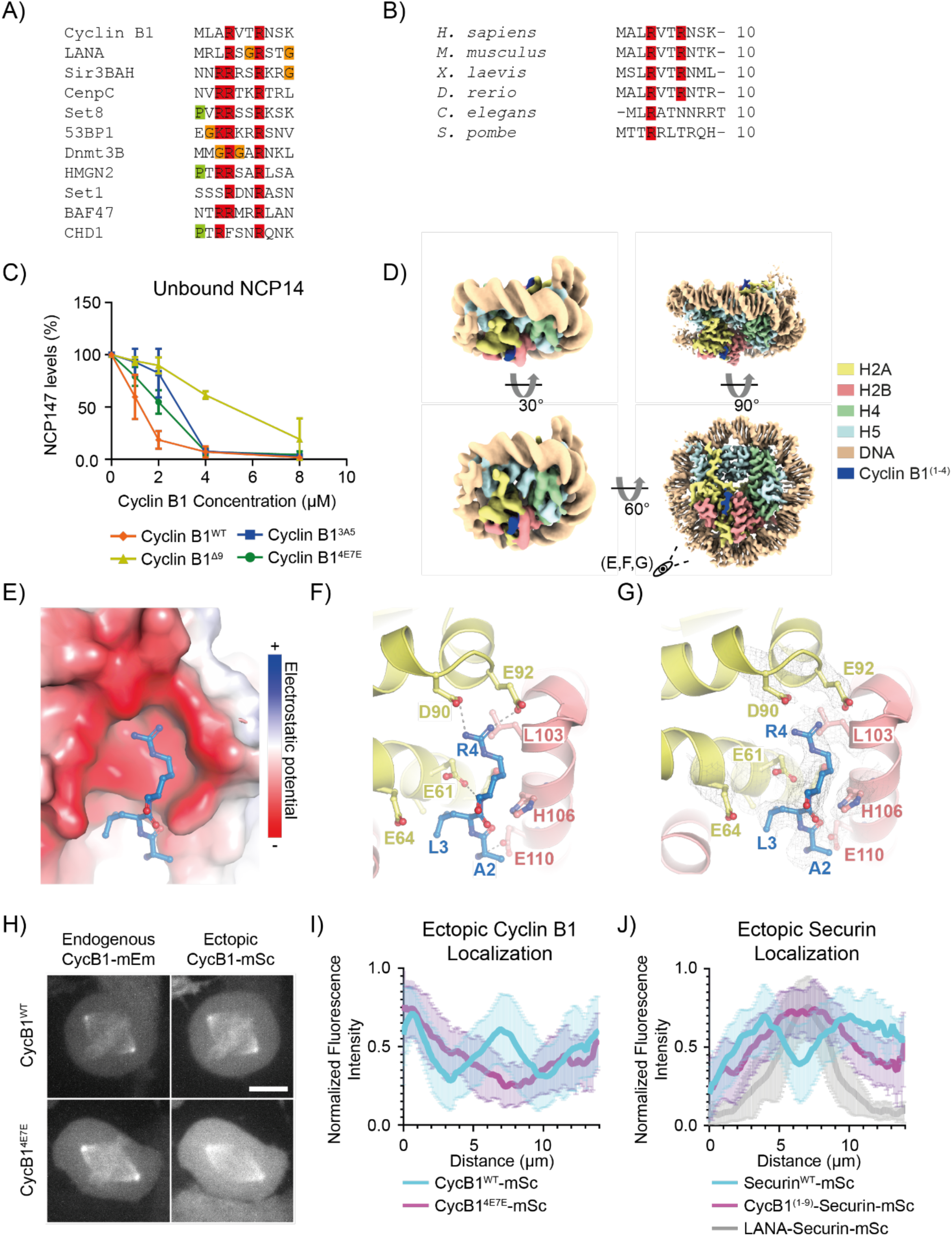
Cyclin B1 N-terminus mediates nucleosome binding. A) Protein alignment of the N-terminus of human Cyclin B1 with known arginine anchors of other nucleosome interacting proteins. B) Protein alignment of the N-terminus of human Cyclin B1 with the same region of Cyclin B1 orthologues. C) Quantification of EMSAs of full-length Cyclin B1 WT and a 4E7E mutant. N = 3 independent experiments. D) Cryo-EM structure of the NCP in complex with Cyclin B N-terminus (NCP^CbNT^). On the left the cryo-EM density is low-pass filtered to 7Å for showing the nucleosome particle in its entirety (including the less rigid entry and exit DNA). On the left the same structure is shown at full power (2.5 Å resolution). All the complex components are coloured as indicated on the right. E). A close-up view of D, residues 2-4 of Cyclin B1 interact with the acidic patch of NCP147. Cyclin B1 is shown as ball and stick, histones H2A and H2B are shown as electrostatic surface potentials (-/+5.000). F) Same structure as in (E), however, H2A and H2B histones are shown as ribbon models. Interacting side-chains are depicted. Dashed lines indicate hydrogen bonds between the arginine 4 of Cyclin B1 and acidic patch residues D90 and E92 Cyclin B1 peptide backbone at A2 interacts with the H2B E110. G) Cryo-EM density on the side-chains shown in (F). H) Maximum projections of confocal images representative of RPE-1 Cyclin B1-mEmerald^+/+^ cells ectopically expressing the indicated variant of Cyclin B1-mScarlet. Scale bar represents 10 μm. I) Line profile graph representing the pixel-by-pixel fluorescence intensity over a line drawn from centrosome to centrosome of RPE-1 Cyclin B1-mEmerald^+/+^ cells ectopically expressing the indicated Cyclin B1 variant. n = 18 cells per condition, N = 3 independent experiments. J) Line profile graph representing the pixel-by-pixel fluorescence intensity over a line drawn from centrosome to centrosome of RPE-1 Cyclin B1-mEmerald^+/+^ cells ectopically expressing the indicated Securin variant. n ≥ 16 cells per condition, N = 3 independent experiments.

To gain further insight into the mode of binding of Cyclin B1 to the nucleosome, we employed cryo-electron microscopy (cryo-EM) to determine the structure of NCP bound to a peptide containing the first 21 residues of Cyclin B1 (NCP^CbNT^). We obtained a reconstruction at 2.5 Å resolution (Table S3), showing residues 2-4 of Cyclin B1 interacting with the nucleosome acidic patch on the NCP (Fig. 4D). Cyclin B1 Arginine 4 establishes hydrogen bonding with the acidic patch residues D90 and E92 in H2A, (Fig. 4E, F, G, S4), and the peptide backbone of Cyclin B1 A2 interacts with H2B E110.

To test whether the N-terminus of Cyclin B1 was required for its chromosomal localisation in living cells, we expressed tetracycline-inducible Cyclin B1-mScarlet variants in the background of our RPE-1 Cyclin B1-mEmerald^+/+^ cells. We reasoned that by comparing the fluorescence intensity of mEmerald and mScarlet we would be able to assess the localisation of ectopically expressed Cyclin B1 using endogenous Cyclin B1 as an internal control (Fig. 4H). To facilitate the analysis, we labelled DNA with siR-DNA and treated cells with the proteasome inhibitor MG132 to induce a metaphase arrest (Potapova et al., 2006). By measuring the fluorescence intensity along a line drawn from centrosome to centrosome, we observed a three-peak pattern for Cyclin B1-mEmerald, whereby the first and the last peaks correspond to the two centrosomes and the middle peak colocalised with the maximum DNA signal (Fig. 4I). Ectopic Cyclin B1^WT^-mScarlet largely overlapped with endogenous Cyclin B1-mEmerald, whereas a Cyclin B1^4E7E^-mScarlet mutant failed to localise to the chromosomes (Fig. 4H, I, Fig S5 A). We obtained comparable results by deleting the first 9 residues of CyclinB1, or by substituting residues 3 to 7 for alanine (Fig S5 A). To determine whether the N-terminus of Cyclin B1 was sufficient to localise a protein to the chromosomes, we tagged Securin with residues 1 to 9 of Cyclin B1. Securin^WT^-mScarlet, was enriched at the mitotic spindle but largely excluded from chromatin (Hagting et al., 2002), but a fusion construct between the Cyclin B1-Securin fusion protein (Cyclin B1^(1-9)^-Securin-mScarlet) displayed partial localisation to chromosomes (Fig. 4J, S5 B, C, D). As a positive control, fusing Securin-mScarlet to a peptide that tightly binds to the nucleosome acidic patch (the LANA (Latency Associated Nuclear Antigen) peptide of Kaposi Sarcoma Herpes Virus, Barbera et al., 2006), strongly enriched Securin on chromosomes (Fig. 4J, Fig S5 C).

**Figure 5.**
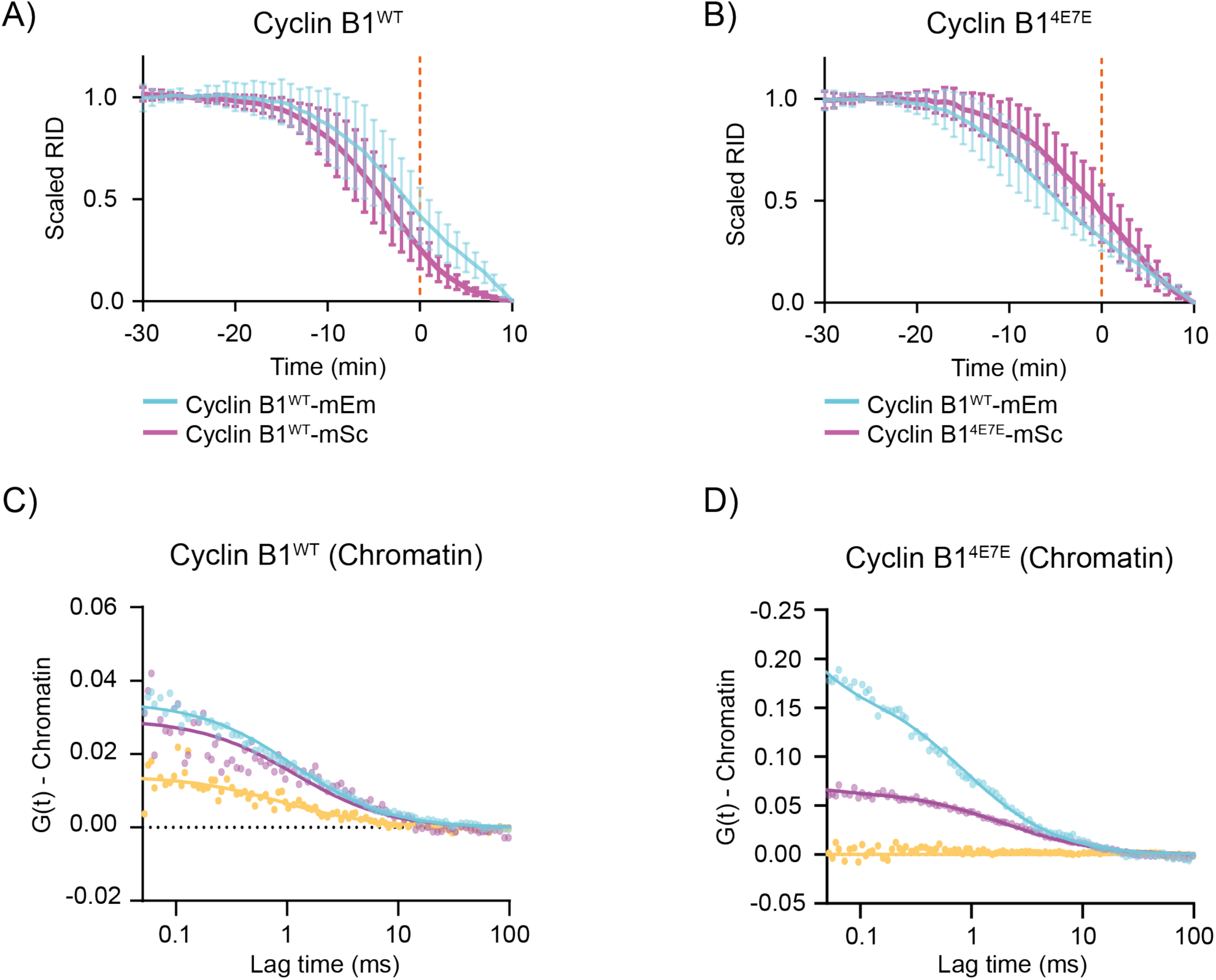
Cyclin B1 nucleosome localization determines its timely degradation. A, B). Plot of the fluorescence intensity of Cyclin B1 over time. n ≥ 17 cells per condition, N ≥ 3 independent experiments. In this and following Cyclin B1 degradation graphs, Cyan: endogenous Cyclin B1-mEmerand, Magenta: ectopically expressed Cyclin B1-mScarlet, unless otherwise specified. C, D) Representative graphs of the autocorrelation of APC8-mScarlet (magenta) and ectopically expressed Cyclin B1-mEmerald (cyan) variant and the cross-correlation (yellow) between the two.

Together, these data demonstrate that the N-terminus of Cyclin B1 acts as an arginine anchor that docks Cyclin B1 to the acidic patch of nucleosomes during mitosis.

### Nucleosome binding is important for timely Cyclin B1 degradation

We previously showed that the first 9 residues of Cyclin B1 were important for timely Cyclin B1 degradation (Matsusaka et al., 2014), providing a potential link between the localisation of Cyclin B1 at chromatin and its degradation. To investigate whether nucleosome binding is required for the timely degradation of Cyclin B1, we compared the degradation of endogenous Cyclin B1-mEmerald with that of ectopically expressed Cyclin B1-mScarlet variants. While Cyclin B1^WT^-mScarlet was degraded slightly before endogenous Cyclin B1, Cyclin B1^3A5^-mScarlet, Cyclin B1^Δ2-9^-mScarlet and Cyclin B1^4E7E^-mScarlet were all degraded later (Fig. 5A, B, S6A). When we measured the interaction between the APC/C and Cyclin B1 using FCCS we found that the cross correlation between ectopically expressed Cyclin B1^WT^-mEmerald and endogenous APC8-mScarlet was stronger on the chromosomes than in the cytoplasm, as expected (Fig. 5C, S6B, C), whereas Cyclin B1^4E7E^-mEmerald only cross-correlated with APC8-mScarlet in the cytoplasm (Fig. 5D, S6C, D). The cytoplasmic cross correlation between Cyclin B1^4E7E^-mEmerald and APC8-mScarlet remained unchanged after treating cells with APCin and proTame, indicating that this interaction was not mediated by the D-box (Fig. S6D).

The delay in the degradation of Cyclin B1 mutants indicated that chromatin localisation might be important for Cyclin B1 degradation, but an alternative hypothesis was that the N-terminus of Cyclin B1 acted as an additional degron to enhance binding to the APC/C. To test this, we assayed Cyclin B1^Δ9^ binding to APC/C using size exclusion chromatography (Fig. 6A, B) and found that purified Cyclin B1 and Cyclin B1^Δ9^ behaved identically when coeluting with the APC/C. This indicated that the arginine-anchor motif of Cyclin B1 did not measurably contribute to APC/C binding.

**Figure 6.**
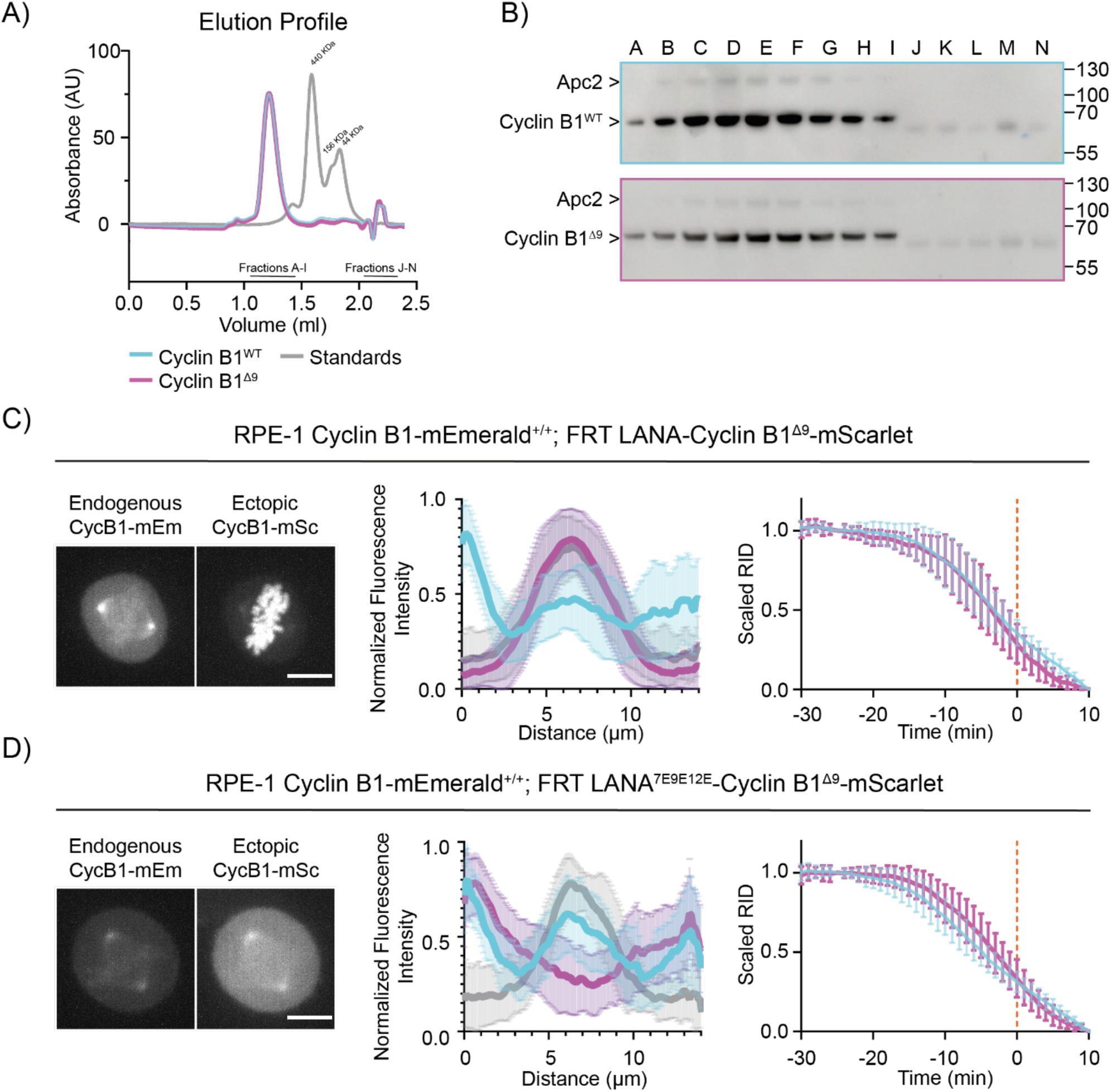
Restoring Cyclin B1^Δ9^ chromatin localisation rescues its degradation timing. A) The elution profile of APC/C^CDC20^ with Cyclin B1^WT^ (cyan) or Cyclin B1^Δ9^ (magenta) when run through a ÄKTA micro. Graph representative of N = 3 independent experiments. B) Representative immunoblot of the size exclusion chromatography showed in A). C, D) Left: Maximum projections of confocal images representative of RPE-1 Cyclin B1-mEmerald^+/+^ cells ectopically expressing the indicated variant of Cyclin B1^Δ9^-mScarlet. Middle: Graphs representing the pixel-by-pixel fluorescence intensity over a line going from centrosome to centrosome of RPE-1 Cyclin B1-mEmerald^+/+^ (Cyan) cells ectopically expressing the indicated variant of Cyclin B1-mScarlet (Magenta). Grey indicates siR-DNA staining. n ≥ 38 cells per condition, N ≥ 3 independent experiments. Right: Cyclin B1 degradation graph representing the fluorescence intensity of Cyclin B1 over time of RPE-1 Cyclin B1-mEmerald^+/+^ (Cyan) cells ectopically expressing the indicated variant of Cyclin B1-mScarlet (Magenta). n ≥ 15 cells per condition, N ≥ 3 independent experiments.

If chromatin localisation is important for Cyclin B1 degradation, we reasoned that restoring the localisation of Cyclin B1^Δ9^ should restore timely degradation. We tested this possibility by fusing the N-terminus of Cyclin B1^Δ9^ to the LANA peptide. In agreement with our hypothesis, the LANA-Cyclin B1^Δ9^-mScarlet fusion protein was highly enriched at chromatin and its degradation rate was similar to that of endogenous Cyclin B1 (Fig. 6C). (Note that it was immaterial whether we fused the LANA peptide to the N or C terminus of the Cyclin (Fig. S7A). Moreover, FCCS measurements confirmed that ectopically expressed LANA-Cyclin B1^Δ9^-mEmerald interacted with endogenous APC8-mScarlet on the chromosomes (Fig. S7B, C). A control fusion between Cyclin B1^Δ9^ and a mutant of LANA unable to bind nucleosomes, failed both to localise to chromatin and to restore Cyclin B1 degradation (Fig. 6D).

Our results raised the question of whether localising Securin, the other metaphase APC/C substrate, to chromatin would be sufficient to enhance its degradation, and indeed we found that ectopically expressed LANA-Securin-mScarlet was highly enriched at chromatin (Fig. 4J) and its degradation began earlier than wild type Securin-mScarlet (Fig. S8A, B). We also observed a slight advance in Securin degradation when it was fused to the first 9 residues of Cyclin B1 (Fig. S8A, B). Overexpressing Securin did not affect the subcellular degradation of Cyclin B1, indicating that competition between APC/C substrates (Kamenz et al., 2015) was not responsible for the delayed degradation of Cyclin B1 in the cytoplasm (Fig. S8C).

We conclude that the binding between the N-terminus of Cyclin B1 and the nucleosome acidic patch confers timely degradation of Cyclin B1.

### CRISPR/Cas9 gene editing to mutate the arginine anchor in Cyclin B1 required simultaneous mutation of p53

Chromosome binding is important for Cyclin B1 degradation to start as soon as the SAC is turned off. We therefore sought to determine the effect on the fidelity of mitosis of eliminating chromosome binding by mutating the arginine anchor in endogenous Cyclin B1.

We used CRISPR/Cas9^D10A^ to introduce the 4E7E mutation into both alleles of Cyclin B1 in RPE1 CCNB1-mEmerand^+/+^ (Fig. 7A). Our initial attempts did not result in any homozygous mutant clones (>2000 clones screened). We did obtain clones when we co-transfected guide RNAs to mutate the tumour suppressor gene TP53 together with those targeting Cyclin B1 (6 potential homozygous clones from 132 colonies screened), as assessed by PCR (Bowden et al., 2020; Chiang et al., 2016). We confirmed homozygous mutations in three independent clones (B1H3, 7A11, 7H10 – hereafter referred collectively to as Cyclin B1^4E7E^ clones) by DNA sequencing (data not shown). Immunoblot analysis revealed that Cyclin B1 levels were considerably reduced in these clones (by ∼ 95% with respect to parental cells, Fig. S9 C, see below). We also generated two RPE1 CCNB1-mEmerand^+/+^; TP53^-/-^ cell lines to use as controls for the effect of knocking out p53 (9B2, 9B3 – hereafter referred collectively to as p53^-/-^ clones), which were confirmed by genomic sequencing (Fig. S9A, B), and by testing for functionality by treating the cells with the p53 stabilising compound Nutlin-3A (Vassilev et al., 2004, Fig. S9C, D – see additional text for Figure S9).

**Figure 7.**
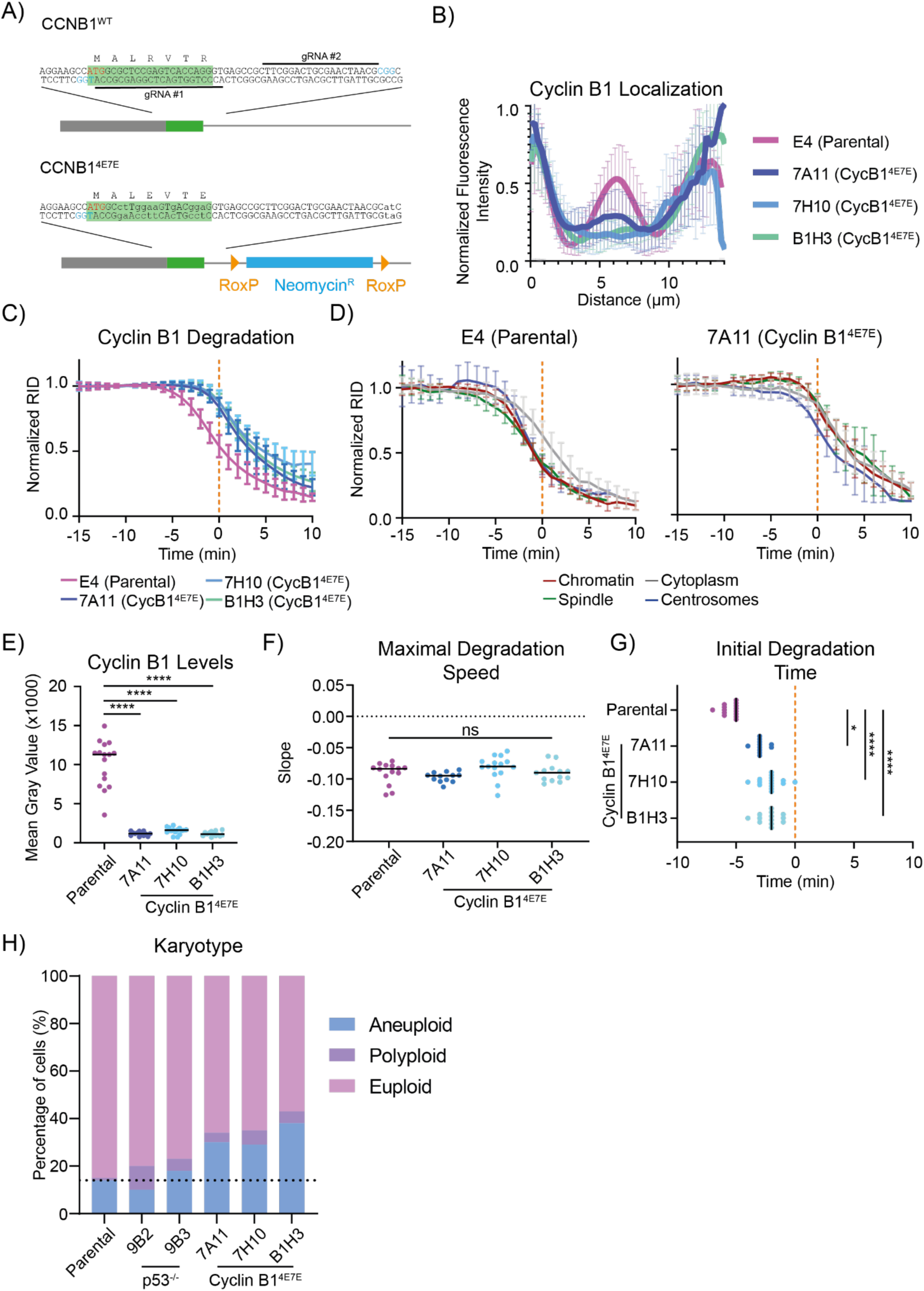
Endogenous Cyclin ^4E7E^ recapitulates ectopic Cyclin 4E7E and increases aneuploidy. A) Schematic representation of the CRISPR strategy used to introduce the 4E7E mutation. The green box indicates CCNB1’s first exon, blue box is a neomycin resistance cassette, grey box indicates the 5’UTR of CCNB1. Yellow triangles indicate RoxP sites. The red text refers to CCNB1’s ATG, blue text refers to PAM sites. B) Graphs representing the pixel-by-pixel **fluorescence** intensity over a line going from centrosome to centrosome of RPE-1 Cyclin B1-mEmerald^+/+^ compared to Cyclin B1^4E7E^ clones. For panels B-G, n ≥ 12 cells per condition, N = 3 independent experiments. C, D) Cyclin B1 degradation graph representing the fluorescence intensity of Cyclin B1 over time of RPE-1 Cyclin B1-mEmerald^+/+^ compared to Cyclin B1^4E7E^ clones. E, F) Dot plots representing the maximal degradation speed (E) or the initial degradation time (F) of RPE-1 Cyclin B1-mEmerald^+/+^ compared to Cyclin B1^4E7E^ clones. Dot plots representing the quantification of the mean fluorescence intensity of Cyclin B1 during metaphase in RPE-1 Cyclin B1-mEmerald^+/+^ compared to Cyclin B1^4E7E^ clones. G) Bar graph representing the karyotype classification from metaphase chromosome spreads of RPE-1 Cyclin B1-mEmerald^+/+^ compared to p53^-/-^ clones and Cyclin B1^4E7E^ clones. Dotted line represents the level of aneuploidy in parental cells.

### The degradation of endogenous Cyclin B1^4E7E^ is delayed and results in aneuploidy

We assayed Cyclin B1 localisation and degradation in RPE-1 CCNB1^4E7E^-mEmerald^+/+^ cells. In all Cyclin B1^4E7E^ clones, line profile analysis of confocal images of metaphase cells showed a strong reduction of Cyclin B1 localisation at chromosomes when compared to parental cell lines (Fig. 7B). Live cell imaging revealed a delay in Cyclin B1 degradation in Cyclin B1^4E7E^ compared to parental Cyclin B1^WT^, both when considering overall levels of Cyclin B1 and at individual subcellular locations (Fig. 7C, D, S10A). In addition, Cyclin B1^4E7E^ did not show any spatial pattern of degradation when comparing the chromosomes, spindle, centrosomes and cytoplasm (Fig. 7D). Note that the loss of p53 alone did not influence Cyclin B1 degradation (Fig. S10B). We obtained comparable result with ectopic expression of Cyclin B1^4E7E^-mScarlet (Fig. S10C). Quantifying the total Cyclin B1 levels in metaphase Cyclin B1^4E7E^ clones, we measured a reduction of over 90% compared to the parental cell line with, confirming our previous western blot result (Fig. 7E, S9C).

To gain further insight into the dynamics of Cyclin B1^4E7E^ degradation, we compared its maximum degradation speed as well as the onset of degradation with those of the parental cell line (see Materials and Methods, Lu et al., 2015). Although the maximum degradation speed did not differ significantly between the Cyclin B1^4E7E^ clones and the parental cell line, our analysis highlighted a significant difference in timing of the onset of degradation (Fig. 7F, G, S10D, E).

Finally, we asked whether the Cyclin B1^4E7E^ mutation had any effect on the fidelity of cell division. Analysis of chromosome spreads showed that 30 to 40% of the cells carrying the Cyclin B1^4E7E^ had become aneuploid compared to only 10 to 15% in the parental cells and p53^-/-^ clones (Fig. 7H).

Altogether these data show that the 4E7E mutation prevents Cyclin B1 localising to chromosomes in mitosis, delays its degradation in metaphase, and cells become more aneuploid.

## DISCUSSION

The destruction of Cyclin B1 is the crucial step that initiates the events leading to chromosome separation and mitotic exit. Twenty years ago, we and others first showed that Cyclin B1 degradation is not homogeneous across the cell: in human cells it disappears earlier from chromosomes, spindle, and centrosomes than from the cytoplasm (Clute and Pines, 1999). Here, we have elucidated the mechanism behind this. We have found that the pattern of Cyclin B1 degradation depends upon it binding to the acidic patch on nucleosomes through its N-terminal arginine anchor, and this cooperates with the Destruction box to enhance Cyclin B1 ubiquitylation by the APC/C and timely degradation in metaphase.

Combining CRISPR/Cas9 gene editing with FCCS has enabled us to measure dissociation constants *in vivo*. Thus, for the first time to our knowledge, we have measured the *in-vivo* dissociation constant between Cyclin B1 and APC/C. This varies between 140 nM at the centrosomes, and 60 nM in the cytoplasm (Table S3). Although these values are likely to be slightly lower than the actual dissociation constant due to technical limitations of FCCS (Barbiero et al., 2022), they are close to the estimated 63 nM obtained *in vitro* using sea urchin Cyclin B1 and fission yeast APC/C (Carroll and Morgan, 2002).

Our conclusion that binding between Cyclin B1 and the APC/C is promoted by nucleosomes is consistent with previous data in human cell lysates showing that APC/C ubiquitination activity toward Cyclin B1 is enhanced in chromatin fractions compared to the cytoplasm (Sivakumar et al., 2014), and that DNA and other polyanions may act as a selectivity barrier facilitating the ubiquitination of D-box-containing substrates over lower-affinity substrates (Mizrak and Morgan, 2019). Our data provide support for the idea that nucleosomes themselves act as a scaffold to bring together APC/C and Cyclin B1. Although APC/C and Cyclin B1 both bind the acidic patch, they are unlikely to compete *in vivo* given the abundance of nucleosomes (∼ 6.23 µM) when compared to the much lower concentrations of both Cyclin B1 and APC/C (about 100 nM, Table S2, (Beck et al., 2011). Our data indicate that nucleosome binding is important for the APC/C to bind productively to Cyclin B1 as soon as the SAC is turned off. We find that a Cyclin B1 mutant that is unable to bind nucleosomes still localises to the spindle and centrosomes but fails to engage APC/C in a D-box dependent manner and its degradation is delayed. Thus, an intriguing possibility is that binding to nucleosomes might expose or optimally present the D-box to the substrate binding site of the APC/C. The D-box itself must have some role in interacting with chromosomes because R42 of the D-box also contributes to chromosome localization (Pfaff and King, 2013), our unpublished observation); for example, it is possible that R42 is important to bridge nucleosome-bound Cyclin B1 with nucleosome-bound APC/C.

An alternative hypothesis to explain the role of chromatin in Cyclin B1 degradation would be that nucleosomes increase the local concentration of Cyclin B1 and APC/C to enhance the kinetics of Cyclin B1 ubiquitylation. Congruent with this idea, we observed that recruiting Securin to the chromatin is sufficient to enhance its degradation and changing the topology of the interaction between Cyclin B1 and nucleosomes (by moving the LANA peptide from N- to C-terminus) has no evident effect on Cyclin B1 degradation.

We do not know whether nucleosome binding is a conserved mechanism to mediate APC/C interaction with Cyclin B1. The N-terminal sequence of Cyclin B1 diverges in non-vertebrates and plants but it is generally arginine-rich, which could mean that nucleosome binding is also conserved. One exception is Drosophila, where the N-terminal arginines of Cyclin B1 are not conserved and it is notable that Drosophila Cyclin B1 first disappears on the spindle poles and only later on chromosomes (Huang and Raff, 1999). Although we did not directly investigate the role of spindle and centrosome localization of Cyclin B1 in its degradation, we found that removing Cyclin B1 from the chromatin delays its degradation throughout the cell, arguing that chromatin is the major site of Cyclin B1 degradation in human cells. This raises the question why Cyclin B1 also appears to be degraded faster at the centrosomes and spindle. A possible explanation is fast diffusion of Cyclin B1 between centrosomes, spindle and chromosomes, such that Cyclin B1 could be rapidly ubiquitylated by a chromosome-bound pool of APC/C and then degraded anywhere between the chromatin and the spindle apparatus. Our FCCS data, however, show that Cyclin B1-APC/C complexes are present at centrosomes and spindle. Thus, we favour an alternative model for Cyclin B1 degradation where nucleosomes promote the formation of a Cyclin B1-APC/C complex that can diffuse to other subcellular locations.

Although an arginine anchor is not present in other cyclins, nor in securin and NEK2A, other APC/C substrates such as KIFC1 (Kinesin-like protein KIFC1), BARD1 (BRCA1-associated RING domain protein 1), TPX2 (Targeting protein for Xklp2), and ZC3HC1 (Zinc finger C3HC-type protein 1), are known to bind the nucleosome acidic patch (Singh et al., 2014; Skrajna et al., 2020; Song and Rape, 2010; Stewart and Fang, 2005; von Klitzing et al., 2011), which raises the possibility that they share an APC/C interaction mechanism similar to that of Cyclin B1. The APC/C also has an important role in G1 phase and in neurodifferentiation, where nucleosome-binding could be important to target chromatin-associated proteins such as Ki67 (Antigen Kiel 67), Top2a (DNA topoisomerase IIα) and the chromosomal passenger complex that accumulate in the cerebellum of APC/C mutant mice and G1 phase of CDH1-mutant RPE1 cells (Ledvin et al., 2023). Multiple mechanisms for chromatin recruitment of the APC/C may exist, however: Oh and colleagues reported that APC/C binding to promoters of ES cells depends on WDR5 to regulate G1 transcription in human embryonic stem cells (Oh et al., 2020).

We found that mutating the arginine anchor of Cyclin B1 results in aneuploidy in otherwise genomically stable RPE-1 cells. That we were not able to introduce the mutation in cells with p53 further indicates that the Cyclin B1 arginine anchor is important for genome stability. Although we cannot formally exclude that aneuploidy originates from lower levels of Cyclin B1 in these cells, it is likely that the reduced levels of Cyclin B1 may be a compensatory mechanism rather than the root cause of aneuploidy. In early mitosis, securin and Cyclin B1 prevent the premature segregation of chromosome by directly binding and inhibiting separase; therefore, lowering the overall levels of Cyclin B1 would guarantee low Cdk1 activity during mitotic exit despite delayed degradation of Cyclin B1. At the end of metaphase, separase must quickly cleave the cohesin ring that holds chromatids together to ensure chromosome segregation is synchronised with mitotic exit. If Cyclin B1 degradation is delayed this will prolong separase inhibition by Cyclin B1, which would result in chromosome segregation defects and consequent aneuploidy.

## MATERIAL AND METHODS

### Cell culture and drug treatment

hTERT RPE-1 FRT/TO cells were cultured in F12/DMEM (Sigma-Aldrich) medium supplemented with GlutaMAX (Invitrogen), 10% FBS (Gibco), 0.35% sodium bicarbonate, penicillin (100 U/ml), streptomycin (100 µg/ml) and Fungizone (0.5 µ/ml). Cells were maintained in a humidified incubator at 37°C and 5% CO2 concentration. For live-cell imaging experiments cells were imaged in Leibovitz L-15 (Thermofisher) medium supplemented with 10% FBS, penicillin (100 U/ml) and streptomycin (100 µg/ml).

In the indicated experiments, cells were stained with 20 nM sirDNA (Spirochrome) following manufacture’s protocol or treated with 100 nM paclitaxel (Sigma-Aldrich), 10 µM MG132 (Selleckchem), 12 µM proTAME (Sigma) and 200 µM APCin (Sigma), Reversine 5 µM (Selleckchem), CenpE inhibitor 50 nM (GSK923295 - Selleckchem), Nutlin-3A (Sigma) 10 µM. Inducible gene expression was performed using 1 µg/ml tetracycline (Calbiochem). In FCCS experiments tetracycline was added 3 h before imaging, in all other experiments tetracycline was added 16 h before imaging. Cells were exposed to Nutlin-3A for 24 h prior to the experiment. Mg132, GSK923295, APCin, proTAME were used for 0.5 - 1 h before the experiment. Reversine and paclitaxel were added immediately before the experiment.

### Gene editing

For APC8 tagging, one million RPE-1 FRT/TO cells or RPE-1 CCNB1-mEmerald^+/+^ FRT/TO cells were transfected using 500 ng of a modified version of the PX466 ‘All-in-One’ plasmid containing Cas9D10A-T2A-mEmerald and gRNAs targeting APC8 (5′-CCACACGCAGAGTTTCTCCA-3′ and 5′-GTCTTCTGTCACGCCATAGT-3′). The all-in-one plasmid was cotransfected with 500 ng of repair plasmid designed as a fusion of GSAGSA-mScarlet flanked by two 500 bp arms, homologous to the genomic region around the Cas9 cutting site. All CRISPR repair templates were cloned into pUC57-Kan plasmid. 72 h post transfection, 50 000 mRuby positive cells were sorted in a 1 cm well and expanded for one week before a second sorting of single cells in 96 well plates. The presence of mScarlet tag was identified through PCR using primers forward 5′-AAGTGGAAAGCCTACCTTGG-3′ and the reverse 5′-GCTGGCTTGAGAGTAGCCAAC-3′. PCR products of positive clones were sequenced using the same primers.

For Cyclin B1^4E7E^ and p53^-/-^ cell lines generation, RPE-1 FRT/TO cells or RPE-1 CCNB1-mEmerald^+/+^ FRT/TO cells were transfected using Cas9^D10A^ RiboNucleoProteins (RNPs – Integrated DNA technology). Briefly, sgRNA were assembled by mixing equal volumes of individual gRNA 100 µM with tracr-RNA 100 µM. For Cyclin B1 the following gRNAs were used: 5’-CCTGGTGACTCGGAGCGCCA-3’ and 5’-CTTCGGACTGCGAACTAACG-3’. For TP53 we used 5’-TCCACTCGGATAAGATGCTG-3’ and 5’-AAATTTGCGTGTGGAGTATT-3’ (Chiang et al. 2016). The mixture was incubated at 95°C for 5 minutes, and then kept at RT for 40 minutes to obtain annealed sgRNAs. 4 uL of annealed sgRNA were then mixed with 2.5 µL of Cas9^D10A^ protein 62 nM to obtain RNPs. Two million cells were transfected with 18.8 pmol of each RNPs (two targeting CCNB1, two targeting TP53), together with 6 µg of a repair plasmid (in the case of Cyclin B1^4E7E^ clones) designed with the wobbled first exon of Cyclin B1 with the 4E7E mutation and a Neomycin resistance cassette, flanked by two 500 bp arms homologous to the genomic region around the Cas9 cutting site. 72 h post transfection, Cyclin B1^4E7E^ cells were selected for 72 h with Nutlin-3A 5 µM and for 7 days with Geneticin (Gibco) 0.4 mg/ml. p53^-/-^ cells were not transfected with any repair template and did not receive the Geneticin selection. Cells were then single cell sorted in 96 well plates. The presence of the 4E7E mutation was identified through PCR and sequencing using primers forward 5′-GCCTTTCATGAACTATATTATTGC-3′ and the reverse 5′-AGCCGCCGCATTGCATCAGC-3′. TP53 status was assessed by PCR and sequencing using primers forward 5′-CTAGTGGGTTGCAGGAGGTG-3′ and the reverse 5′-TAAGCAGCAGGAGAAAGCCC-3′. To distinguish the two alleles of TP53 PCR product were subjected to cloning into TOPO2.1 plasmid prior to sequencing, according to the manufacture’s protocol.

For Cyclin B1-mScarlet, Securin-mScarlet and Cyclin B1-mEmerald variants ectopic expression, RPE-1 FRT/TO CCNB1-mEmerald^+/+^, RPE-1 FRT/TO CCNB1-mEmerald^+/+^; APC8-mScarlet^+/+^ or RPE-1 FRT/TO APC8-mScarlet^+/+^ cell lines were cotransfected with the relevant gene CDS cloned into pcDNA5-FRT/TO (p1795) and pOG44 (Invitrogen) using a 1:5 ratio (using 1 µg total DNA per million cells). All transfections were followed by a two-week selection using Geneticin (Gibco) 0.4 mg/ml. Cell bearing Cyclin B1-mScarlet variants were sorted into low and high expressing, only low expressing cells were selected for further analysis. Cells expressing Cyclin B1^R42AL45A^-mEmerald mutant were not sorted due to the cytoxicity of the construct.

All transfections were performed by electroporation using a Neon Transfection System (Invitrogen) with two pulses at 1400 V for 20 ms.

### Protein extraction and immunodepletion

Cells were trypsinised and incubated in lysis buffer (150 mM NaCl, 50 mM Tris pH 7.4, 0.5% NP-40) supplemented with HALT protease/phosphatase inhibitor cocktail (Thermo Fisher Scientific) for 30 min at 4°C before clarification. Lysates were then clarified through centrifugation (14′000g, 20 min, 4°C) and then quantified using Bradford Reagent (Bio-Rad Laboratories) according to manufacturer’s instructions.

For APC4 immunodepletion, 200 µg of clarified lysates were diluted to a final concentration of 2 µg/µl and incubated with 30 µl of Dynabeads prebound to 3.75 µg of either anti-APC4 Antibody (Movarian Biotech) or Mouse Ig-G, in a total volume of 100 µl for 2 h at 4°C. 40 µg of either the immunodepleted lysates or the input were used for SDS-PAGE.

### Immunoblotting

Cell lysates were separated through SDS-PAGE on a 4–12% NuPAGE gel (Invitrogen) and transferred to an Immobilon-FL polyvinylidene fluoride membrane (IPFL00010, Millipore). The membrane was blocked with 5% Milk, 0.1% Tween, PBS and incubated overnight with primary antibodies at 4°C in 2.5% Milk, 0.1% Tween, PBS. The following day the membrane was washed with 0.1% Tween PBS and incubated with secondary antibodies in 2.5% Milk, 0.1% Tween, PBS for 1 h at RT. Membranes were visualised with LI-COR Odyssey CLx scanner (LI-COR Biosciences). Western blot quantification in figure S1F was performed calculating the area under the curve using Fiji’s ‘Gel’ plugin. Values were adjusted by fitting to a straight-line function obtained by immunoblotting serial dilutions of protein lysate.

In EMSA assays and size exclusion chromatography membranes were blocked overnight in 5 % BSA. Western blot was developed using the ImageQuant 800 (Amersham).

Primary antibodies were used at the indicated concentrations: anti-APC2 (1:1000, Movarian Biotech), anti-APC4 (1:1000, Movarian Biotech), anti-APC8 (1:1000, D5O2D – Cell Signalling), Strep-MAB (1:10000, IBA 2-1507-001 - IBA), anti-p53 (1:1000, DO-1 – Santa Cruz), anti-p21 (1:1000, 12D1 – Cell Signalling), anti-β Tubulin (1:10000, Ab6046 - Abcam), anti-Cyclin B1 (1:1000, GNS1 - Santa Cruz).

Secondary antibodies were: IRDye800CW donkey anti-mouse (926-32212, LI-COR), IRDye800CW donkey anti-rabbit (926-32213, LI-COR), IRDye680CW donkey anti-mouse (926-68072, LI-COR), and IRDye680CW donkey anti-rabbit (926–68073, LI-COR), HRP-conjugated goat anti–mouse (Amersham), HRP-conjugated goat anti–rabbit (Amersham). Secondary antibodies were all used at 1:10’000.

### Expression of proteins in insect cells

Codon optimised cDNA of wild-type and mutant Cyclin B1-StrepII were ordered in pFastBac1 vectors from GeneArt (Thermo Fischer Scientific) and transformed into MultiBac cells for insect cell expression. Sf9 cells were used to create the baculoviruses which were then used to infect High Five cells at a cell density of 1.5X106 cells/mL. High Five cells were incubated for 72 hours at 27 °C, 130 rpm.

To express the APC/C three separate baculoviruses were used to infect High Five cells: the first contained APC/C subunits Apc5, APC8, Apc10, Apc13, Apc15, Apc2 and Apc4-strepII, gifted by David Barford’s lab. The second contained Apc11 and Apc1 phosphomimic mutant (Apc1 Ser364Glu, Ser372Glu, Ser373Glu, Ser377Glu; Zang et al. 2016). The final baculovirus contained subunits Apc3, Apc6, Apc7, Apc12 and Apc16. A cell density of 2X106 was infected with these baculoviruses and then incubated for 72 hours at 27 °C, 130 rpm.

### Protein Purification

All purification steps were carried out at 4 °C:

#### APC/C

Cell pellets were thawed on ice in wash buffer (50 mM HEPEs pH8.3, 250 mM NaCl, 5 % glycerol, 2 mM DTT, 1 mM EDTA and 2 mM Benzamidine supplemented with 0.1 mM PMSF, 5 units/mL Benzonase and an EDTA-free protease inhibitor (Roche). After sonication, the cells were centrifuged for 1 hour at 48,000 xg. The supernatant was bound to a 3X5 mL StrepTactin Superflow Plus cartridges (Qiagen) using a 1 mL/min flow rate. The column was washed extensively with APC/C wash buffer and eluted with wash buffer containing 2.5 mM desthiobiotin (IBA-Lifesciences). Fractions containing APC/C were incubated with tobacco etch virus (TEV) protease overnight. The sample was prepared for anion exchange chromatography by a two-fold dilution with saltless Buffer A (20 mM HEPEs pH 8.0, 125 mM NaCl, 5 % glycerol, 2 mM DTT, 1 mm EDTA). The sample was then loaded onto a 6 mL ResourceQ anion-exchange column (GE Healthcare), and the column washed extensively with buffer A. An elution gradient with Buffer B (20 mM HEPEs pH 8.0, 1 M NaCl, 5 % glycerol, 2 mM DTT, 1 mM EDTA) was used to elute the APC/C. The APC/C was concentrated and the centrifuged (Optima TLX Ultracentrifuge) at 40,000 rpm for 30 minutes.

#### CDC20

CDC20 was purified as previously described (Zhang et al., 2016).

#### Cyclin B1

Wild-type and mutant cyclin B1 cell pellets were thawed in Cyclin B1 wash buffer (50 mM HEPEs pH 8.0, 500 mM NaCl, 5 % glycerol and 0.5 mM TCEP supplemented with 5 Units/mL Benzonase and an EDTA-free protease inhibitor cocktail (Roche)). Cells were sonicated and centrifuged at 55,000 x g for 1 hour. The supernatant was loaded onto a 5 mL StrepTactin Superflow Plus catridge (Qiagen) using a 1 mL/min flow rate and washed with Cyclin B1 wash buffer and eluted with wash buffer containing 2 mM desthiobiotin. Cyclin B1 was concentrated and loaded onto a HiLoad 16/600 Superdex 200 pg column (Cytiva).

### Peptide preparation

A peptide of the first 21 residues of human Cyclin b1, MALRVTRNSKINAENKAKINM, (NeoBiotech) was dissolved in water.

The first 23 residues of Kaposi’s sarcoma-associated herpesvirus, latency associated nuclear antigen (LANA) protein, MAPPGMRLRSGRSTGAPLTRGSC (GenScript) was dissolved in water.

### NCP147 assembly

Recombinant H2A, H2B, H3 and H4 were expressed in *Escherichia coli*, all histones were from Homo sapiens apart for H2B which was from *Xenopus laevis*. The histones were purified and assembled with Widom 601 147bp DNA sequence as previously described (Luger et al., 1999).

### Cryo-EM grid preparation

Quantifoil R1.2/1,3 Cu 300 grids were glow discharged using 15 mA for 1 minute on each side of the grid using an Easiglow (Pelco).1.68 µM of NCP was mixed with 411 µM of CyclinB1^(1-21)^ peptide in 20 mM HEPES pH 7.5, 20 mM NaCl, 0.1 mM TCEP. 2 µL of the premixed sample was added to the grid. After a 5 second wait the grids were blotted for 5 seconds with a blot force of 3, at 4°C, 100 % humidity. The grids were frozen in liquid ethane using a Vitrobot mark IV (Thermo Fisher).

### Cryo-EM data collection and processing

We collected 4,589 EER images of the NCP^CbNT^ sample at ICR on a Glacios Cryo-TEM 200 kV equipped with a Falcon4i operated in counting mode at a pixel size of 0.567 Å per pixel. Movie stacks were frame aligned and binned 4 times. Images with resolution better than 6 Å and a total motion of 30 pixels (estimated during frame alignment) were selected for further processing. A blob picker was used for initial particle picking during the live processing, and for obtaining initial 2D class averages. Templates generated from the latter, were used for template picking and TOPAZ (Bepler et al., 2019; Punjani et al., 2017) training and picking with cryoSPARC. Selected particles from 2D classifications performed with particles picked with the different methods explained above were pooled together and duplicated particles were removed by using the “remove duplicates” function implemented in cryoSPARC. This step removed doubly picked particles within 100 Å (the length of the NCP is ∼100 Å). These particles were piped into the “ab initio model” function implemented in cryoSPARC. These particles were then imported in RELION-3.1.1 (Zivanov et al. 2020) by using pyem and the csparc2star.py script by Daniel Asarnow: (Daniel Asarnow, Eugene Palovcak, & Yifan Cheng. (2019). asarnow/pyem: UCSF pyem v0.5 (v0.5). Zenodo. https://doi.org/10.5281/zenodo.3576630) for 3D refinements. Map improvement was performed by performing sequential 3D classification without alignment (T=4, T=40 and T=80) with a soft mask coupled with 3D refinements, Bayesian polishing steps and Ctf refinements in RELION. This yielded a map at the resolution of 2.5 Å (Fig. S4).

The PDB ID:3LZ0 was used as initial template for 30 building in Coot (Casañal et al., 2020). Cyclin B N-terminus was built de novo based on the excellent quality of our cryo-EM map. Structural model refinement was performed with PHENIX real-space refinement (Afonine et al., 2018) at the resolution of 2.5 Å.

### Size Exclusion Chromatography

Samples were run through a Superose 6 Increase 3.2/300 2.4 mL column on the ÄKTAmicro system in 20 mM HEPES pH 8.0, 150 mM NaCl, 0.5 mM TCEP in a 30 µL reaction volume. 1.5 µM APC/C, 3.3 µM CDC20 and 3.3 µM of Cyclin B1 were mixed and placed on ice for 5 minutes before injection. When running the Cyclin B1 constructs alone 18 µM was injected. Selected fractions were run on SDS-PAGE then transferred to a PDVF membrane.

### Electrophoretic mobility shift assay

For EMSAs of the NCP with cyclinB1, 0.5 µM of NCP147 was mixed with full length cyclin B1 or the n-terminal mutants at a ratio of 1:0, 1:2, 1:4, 1:8 and 1:16 in a 6 µL reaction volume. The reaction buffer used was 20 mM HEPEs pH 7.5, 50 mM NaCl, 0.1 mM TCEP. Reactions were incubated on ice for 10 minutes before adding 5 % (v/v) sucrose. The samples were run on a 5% polyacrylamide gel in 0.25X TBE running buffer at 100 V for 90 minutes. The gels were then stained with SYBR safe and scanned using a Typhoon FLA 9500 imager.

Quantification was done using ImageJ and the band intensities normalised compared to the band intensity of the NCP without protein complex. For EMSAs with the APC/C-CDC20-Cyclin B1^NTD^, strep tagged APC/C was pre-incubated on ice with streptavidin conjugated Alexa Fluor 700 (ThermoFisher Scientific S21383) for 30 minutes. This was then mixed with CDC20 and unlabelled Cyclin B1^NTD^ at a 1:1:1 molar ratio. 0.2 µM NCP147 was mixed with this APC/C solution at ratios of 1:0, 1:1, 1:2, 1:4, 1:8 and 0:1 in 6 µL volumes. The reactions were treated as with the cyclinB1 EMSAs and then run on a 1% agarose gel made in 0.5X TBE and run in a 0.25X TBE running buffer. This was run for 90 minutes at 100 V.

### XL-MS

1.4 µM of APC/C, CDC20 and Cyclin BNTD were mixed to give a 1:1:1 molar ratio. 0.43 µM NCP was mixed with 3.4 µM of the APC/C-CDC20-Cyclin B1^NTD^ mix and incubated on ice for 10 minutes. 1.44 mM of DSSO (disuccinimidyl sulfoxide, A33545, Thermo Scientific) dissolved in DMSO was added to the proteins to give a final volume of 110 µL, this was incubated on ice for 1.5 hours. The sample was quenched with 50 mM TRIS-HCl pH 8.0. After the crosslinking reaction, triethylammonium bicarbonate buffer (TEAB) was added to the sample at a final concentration of 100 mM. Proteins were reduced and alkylated with 5 mM tris-2-carboxyethyl phosphine (TCEP) and 10 mM iodoacetamide (IAA) simultaneously for 60 min in dark and were digested overnight with trypsin at final concentration 50 ng/μL (Pierce). Sample was dried and peptides were fractionated with high-pH Reversed-Phase (RP) chromatography using the XBridge C18 column (1.0 × 100 mm, 3.5 μm, Waters) on an UltiMate 3000 HPLC system. Mobile phase A was 0.1% v/v ammonium hydroxide and mobile phase B was acetonitrile, 0.1% v/v ammonium hydroxide. The peptides were fractionated at 70 μL/min with the following gradient: 5 minutes at 5% B, up to 15% B in 3 min, for 32 min gradient to 40% B, gradient to 90% B in 5 min, isocratic for 5 minutes and re-equilibration to 5% B. Fractions were collected every 100 sec, SpeedVac dried and pooled into 12 samples for MS analysis.

LC-MS analysis was performed on an UltiMate 3000 UHPLC system coupled with the Orbitrap Ascend mass spectrometer (Thermo Scientific). Each peptide fraction was reconstituted in 30 μL 0.1% TFA and 15 μL were loaded to the Acclaim PepMap 100, 100 μm × 2 cm C18, 5 μm trapping column at 10 μL/min flow rate of 0.1% TFA loading buffer. Peptides were then subjected to a gradient elution on a 25 cm capillary column (Waters, nanoE MZ PST BEH130 C18, 1.7 μm, 75 μm × 250 mm) connected to the EASY-Spray source at 45 °C with an EASY-Spray emitter (Thermo, ES991). Mobile phase A was 0.1% formic acid and mobile phase B was 80% acetonitrile, 0.1% formic acid. The gradient separation method at flow rate 300 nL/min was as follows: for 80 min gradient from 5%-35% B, for 5 min up to 95% B, for 5 min isocratic at 95% B, re-equilibration to 5% B in 5 min, for 5 min isocratic at 5% B. Precursors between 380-1,400 m/z and charge states 3-8 were selected at 120,000 resolution in the top speed mode in 3 sec and were isolated for stepped HCD fragmentation (collision energies % = 21, 27, 34) with quadrupole isolation width 1.6 Th, Orbitrap detection with 30,000 resolution and 70 ms Maximum Injection Time. Targeted MS precursors were dynamically excluded for further isolation and activation for 45 seconds with 10 ppm mass tolerance.

Identification of crosslinked peptides was performed in Proteome Discoverer 3 (Thermo) with the MS Annika search engine node (Pirklbauer et al. 2021) for DSSO / +158.004 Da (K). Precursor and fragment mass tolerances were 10 ppm and 0.02 Da respectively with maximum 4 trypsin missed cleavages allowed. Carbamidomethyl at C was selected as static modification and oxidation of M as dynamic modification. Spectra were searched against a FASTA file containing the sequences of the proteins in the complex concatenated with 1000 random UniProt E. Coli sequences as negative control. Crosslinked peptides were filtered at FDR<0.01 separately for intra/inter-links using target-decoy database search. XL-MS data was visualised with xiview.

### Live-cell imaging & image analysis

Mitotic time measurements were obtained using differential interference contrast (DIC) imaging on a Nikon Eclipse microscope (Nikon) equipped with a 20 × 0.75 NA objective (Nikon), a Flash 4.0 CMOS camera (Hamamatsu) and an analyser in the emission wheel for DIC imaging. Single plane images were taken every 3 min, for 24 h using micromanager software (µManager). Where indicated, cells received a treatment with paclitaxel.

Images and quantifications of Cyclin B1 and Securin, excluding FCS and FCCS (see below) were obtained on a Marianas confocal spinning-disk microscope system (Intelligent Imaging Innovations, Inc.) equipped with a laser stack for 445 nm/488 nm/514 nm/561 nm lasers, a 63 × 1.2 NA objective (Carl Zeiss), a Flash4 CMOS camera (Hamamatsu) and Slidebook 6 software (Intelligent Imaging Innovation, Inc.).

For figure 1, 8 Z stacks (Step size = 1 µm) were taken every 30 s, for 90 min, using 20% 488 nm laser power and 10% 647 nm laser power, for 50 ms exposure, 1x1 binning. Image stacks were maximum projected over z-axis and mean fluorescence intensity at different subcellular location was quantified. After background subtraction, each measurement was normalised on the value at 20 frames prior to anaphase and corrected for bleaching using the measurement obtained in the same subcellular location in a sample treated with 50μg/ml cycloheximide (Merk) and 10 µM MG132. Where indicated, cells received a treatment with MG132 or APCin and Tame for 30 minutes prior to filming.

For figure 2, cells were exposed for 30 minutes to 50 nM CenpE inhibitor and treated with 5 uM Reversine immediately before imaging. 10 Z stacks (Step size = 1 µm) were taken every 30 seconds, using 7% 488 nm for 150 ms, and 5% 647 nm laser power, for 100 ms exposure, 4x4 binning. After background subtraction, a region of interest containing the chromatids of a polar chromosomes was manually selected. In such region Cyclin B1 signal was quantified inside and outside the DNA area. Each measurement was normalised on the value obtained at 25 frames prior to anaphase.

For Cyclin B1 and Securin line profile assay, cells were exposed to MG132 for 30 minutes prior to filming. 51 Z stacks (Step size = 0.2 µm) were taken for single time point, using 10% 488 nm, 15% 561 nm and 5% 647 nm laser power, for 100 ms exposure, 1x1 binning. Image stacks were summed over z-axis and the fluorescence intensity over a 10-pixel thick line going from centrosome to centrosome was quantified. After background subtraction, each measurement was scaled between the maximum and minimum value, set as 1 and 0, respectively. In case of Cyclin B1^4E7E^ clones the line-profile was obtained on the same image used to quantify Cyclin B1 degradation (see below), 10 minutes before anaphase.

For Cyclin B1-mScarlet and Securin-mScarlet degradation assay, 10 Z stacks (Step size = 1 µm) were taken every minute, using 10% 488 nm, 10% 561 nm and 10% 647 nm laser power, for 50 ms exposure, 2x2 binning. Image stacks were maximum projected over z-axis and raw integrated density of the whole cell or of specific subcellular locations was measured. After background subtraction, each measurement was either scaled between the value at 25 frames prior to anaphase and the minimum value, arbitrarily set as 1 and 0, or normalised to the value at 25 frames prior to anaphase.

In the quantification of Cyclin B1 degradation in Cyclin B1^4E7E^ clones, Cyclin B1 levels were normalised to frame -10 prior to anaphase and bleaching-corrected to a straight-line function fitted between frame -30 and -25 prior to anaphase. To obtain the maximum degradation speed and time, we fitted a straight-line function using 5 points around the minimum of the first derivative of Cyclin B1 degradation data smoothed with a 4-values averaging window. Maximum degradation speed is the slope of that function. To find the initial degradation point, data were smoothed with a 4-values averaging window and then fitted to a straight-line function using a sliding 5-values window. The initial degradation point is the initial value of the first 5-value window whose slope is below -0.05. Initial degradation speed is the slope of such window.

All image processing and analysis was performed using Fiji (ImageJ).

### Fluorescence Correlation and Cross-Correlation Spectroscopy

A Leica TCS SP8 confocal microscope (DMI8; Leica) integrated with wavelength-adjustable pulsed white light laser and Leica HyD SMD (single molecule detection) detectors was used for all FCS and FCCS measurements. The samples were imaged and measured using a Leica HC PL APO CS2 63x/1.20 water immersion with a pinhole size of 1 airy unit. For proteins tagged with mEmerald, the wavelength was set at 488 nm with a detection range of 505–540 nm, while the wavelength was set at 569 nm and detection range at 580–625 nm for measuring proteins tagged with mEmerald.

Calibration of the instrument prior to each FCS and FCCS experiment and FCCS control experiments (positive and negative) were performed as described in Barbiero et al. 2022. The auto- and cross-correlation functions were fitted using Leica LAS X SMD FCS module. Selection of fitting models, determination of diffusion coefficients, calculation of cross-correlation quotient (q) and dissociation constant (KD) were also performed as discussed in (Barbiero et al., 2022).

### Chromosome spreads

For chromosome spreads, following a 3-hours treatment with 100 ng ml−1 colcemid (GIBCO) cells were trypsinized and recovered in a falcon tube. Cell suspension was centrifuged for 3 min at 250 g and resuspended in 5 ml of 75 mM KCl, added dropwise. After a 15 minute-incubation at 37°C, 10 drops of Carnoys Fixative (3 : 1 methanol : acetic acid) were added. Following a 5 min centrifugation at 200 g, the cell pellet was resuspended in 5 ml of Carnoys Fixative. After 90 min at −20°C, a second fixation was performed using 5 ml of Carnoys Fixative at room temperature for 15 min. Cells were then centrifuged at 200 g for 5 min, the supernatant removed, and the pellet resuspended in 200 µl of Carnoys Fixative. Spreads were performed by dropping the cells on wet slides in a wet chamber from 30 to 40 cm height. Spreads were mounted and stained using Vectashied with DAPI (ReactoLab). Samples were imaged on a GE wide-field DeltaVision Elite microscope equipped with an 63x/1.2-NA Oil Objective, a ImageXpress Micro widefield system camera with appropriate filters and SoftWoRx Imaging software. The number of chromosomes per cell was counted using ImageJ software.

### Protein alignment

For protein alignment displayed in Fig. 2A, we extracted the arginine anchors of the following proteins: LANA-1 (uniprot Q9QR71), Sir3 BAH (P06701), CenpC (Q03188), Set8 (Q9NQR1), 53BP1 (Q12888), DNM3B (Q9UBC3), HMGN2 (P05204), Set1 (P38827), BAF (Q12824) and CHD1 (P32657). For alignment in Fig 2B we used full-length Cyclin B sequences from different species: *H. sapiens* (P14635), *M. musculus* (P24860), *X. laevis* (P13350), *D. rerio* (Q9IB44), *C. elegans* (Q10653) and *S. Pombe* (P10815). For the sequence alignment of the APC3^loop^, the following homologous sequences were taken from ProViz: *H. sapiens* (P30260), *B. taurus* (A7Z061), *A. thaliana* (A0A178VWS3), *D. melanogaster* (Q9VS37), *G. gallus* (Q5ZK91), *X. tropicalis* (Q0P4V8), *C. elegans* (Q9N593), *S. cerevisiae* (P38042), *S. pombe* (P10505). All alignments were performed using MUSCLE (EMBL-EBI) with default settings and colour coded matching the Clustal colour scheme.

### Alphafold

alphafold/2.3.0 was used with the multimer option to predict the NCP:APC3 binding regions (Evans et al., 2022). The input sequences included APC3364-387 and the histone octamer proteins.

### Statistics & Figure assembly

Details on statistics are summarised in Table S1. Data analysis was performed with Python 3.7.0. Statistical analysis and plotting were performed with Prism 8 (GraphPad). Mitotic time graphs were realised following the ‘Superplots’ pipeline (Lord et al., 2020). Figures were assembled using Adobe Illustrator (Adobe).

## ACKNOWLEDGMENTS

We wish to thank Fay Cooke and Anja Hagting, for the early work on the spatial regulation of Cyclin B1 degradation, and Martina Barbiero for the initial FCS data on Cyclin B1. We thank Prof. Stephen Jackson for sharing the RPE-1 TP53^-/-^ cell line. We acknowledge Jing Yang and Ziguo Zhang from David Barford’s laboratory for their help with the APC/C baculovirus generation.

We thank all the members of the Pines and Alfieri groups for helpful discussions. We gratefully acknowledge the support of the ICR Core facilities, in particular, light microscopy and flow cytometry.

L.C., S.V., C.C. and J.P. were supported by an Investigator Award from Wellcome (209470/Z/17/Z), R.Y., R. M. and C.A. were supported by the Sir Henry Dale Fellowship 215458/Z/19/Z. R.Y and R.M. were also supported by the Institute of Cancer Research (ICR - GFR005X, GFR146X). C.P. was funded by a internship from Bologna University, M.M. was supported by a BSBC summer student grant, A.A. was supported by the In2Research program.

## AUTHORS’ CONTRIBUTION

L.C.: investigation, data analysis, writing—original draft, conceptualization, funding acquisition; R.Y.: investigation, data analysis, writing—original draft; S.V.: investigation, data analysis; A,R.: investigation, data analysis; M.M: investigation, data analysis; A,A.: investigation, data analysis; C.P.: investigation, data analysis; C.C.: investigation; R. M.: investigation; T. R.: investigation; J. C.: funding acquisition; C.A.: investigation, conceptualization, data curation, funding acquisition, project administration, writing—original draft. J. P.: conceptualization, data analysis, funding acquisition, project administration, writing—original draft.

## CONFLICT OF INTEREST DECLARATION

The authors declare no conflicts of interest.

## SUPPLEMENTARY MATERIALS

**Figure S1.**
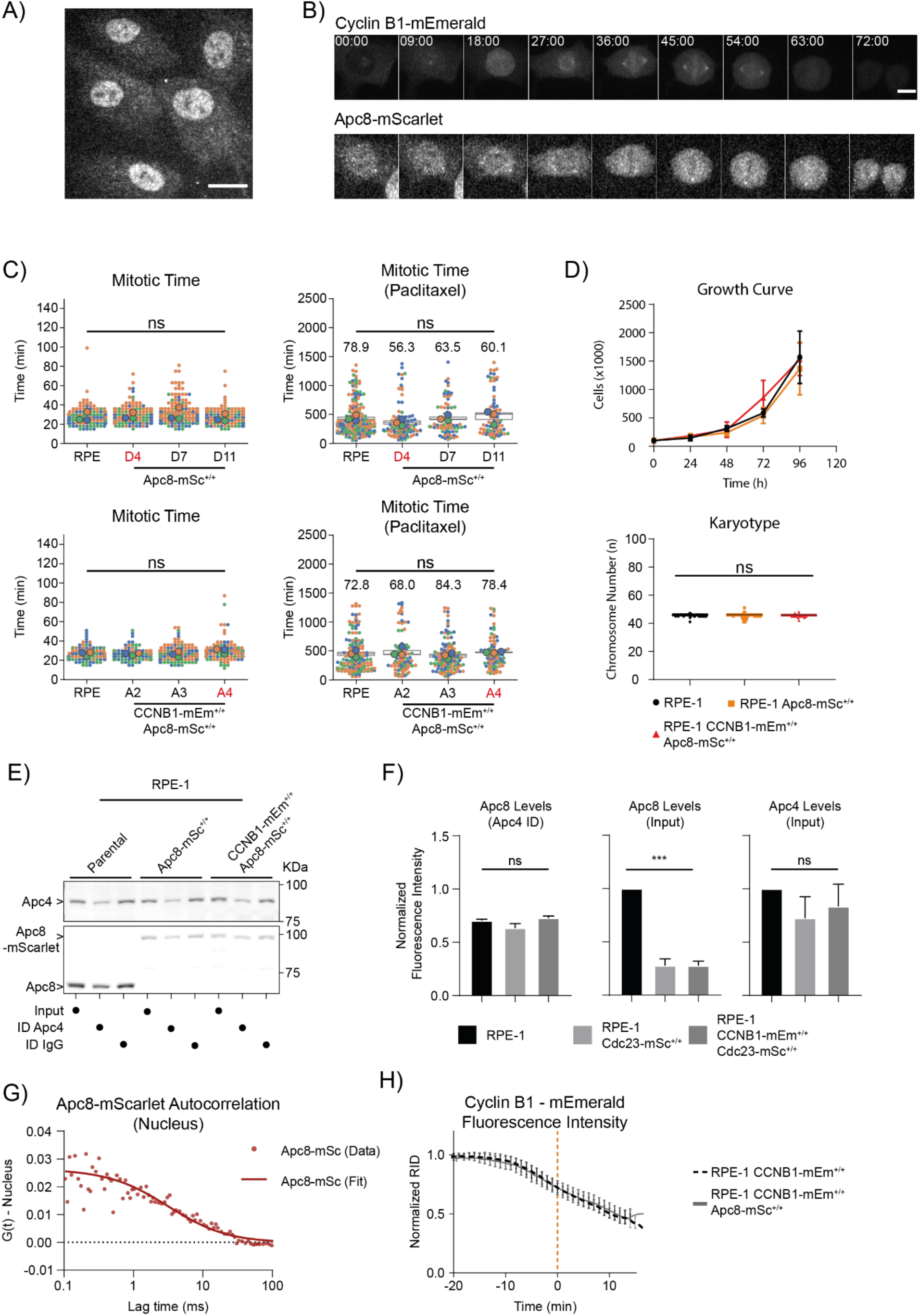
Characterization of RPE-1 CCNB1-mEmerald^+/+^ cells. A) Representative fluorescence confocal image of RPE-1 CCNB1-mEmerald^+/+^; APC8^+/+^ cells in interphase. Scale bar corresponds to 20 μm. B) Representative fluorescence confocal images over time of a CCNB1-mEmerald^+/+^; APC8^+/+^ cells progressing through mitosis. Time is expressed as mm:ss. Scale bar correspond to 10 μm. C) Dot plots of the mitotic timing of parental RPE-1, RPE-1 APC8-mScarlet^+/+^, RPE-1 CCNB1-mEmerald^+/+^ and APC8-mScarlet^+/+^ cells, untreated (left) or treated with 100 nM paclitaxel (right). Each small dot represents one cell, large dots represent the median of independent experiments. n ≥ 83 cells per condition, N = 3 independent experiments. Numbers on the graphs indicate the percentage of cells completing mitosis during the time of observation. Clones marked in red are the ones selected for all following experiments. D) Top, growth curve of RPE-1, RPE-1 APC8-mScarlet^+/+^, RPE-1 CCNB1-mEmerald^+/+^, N = 3 experiment. Bottom, Dot plot of the chromosome number of parental RPE-1, RPE-1 APC8-mScarlet +/+, RPE-1 CCNB1-mEmerald+/+ and APC8-mScarlet^+/+^ cells. Each dot represents chromosome spread. n ≥ 34 spreads per condition, N = 1 experiment. E) Representative Anti-APC8 and anti-Apc4 immunoblot of cell lysates from parental RPE-1, RPE-1 APC8-mScarlet^+/+^, RPE-1 CCNB1-mEmerald^+/+^ and APC8-mScarlet^+/+^ cells before and after immunodepleting Apc4, compared with control immunodepletion with IgG. F) Bar graph representing the quantification of immunoblot in panel E. N = 2 independent experiments. G) Graph representing the autocorrelation function of APC8-mScarlet over time in the nucleus. H) Quantification of normalised Cyclin B1 fluorescence levels over time measured by spinning disk fluorescence microscopy in RPE-1 CCNB1-mEmerald^+/+^ cells compared to RPE-1 CCNB1-mEmerald^+/+^; RPE-1 APC8-mScarlet^+/+^, RPE-1 CCNB1-mEmerald^+/+^ cells. n ≥ 9 cells per condition, N = 3 independent experiments.

**Figure S2.**
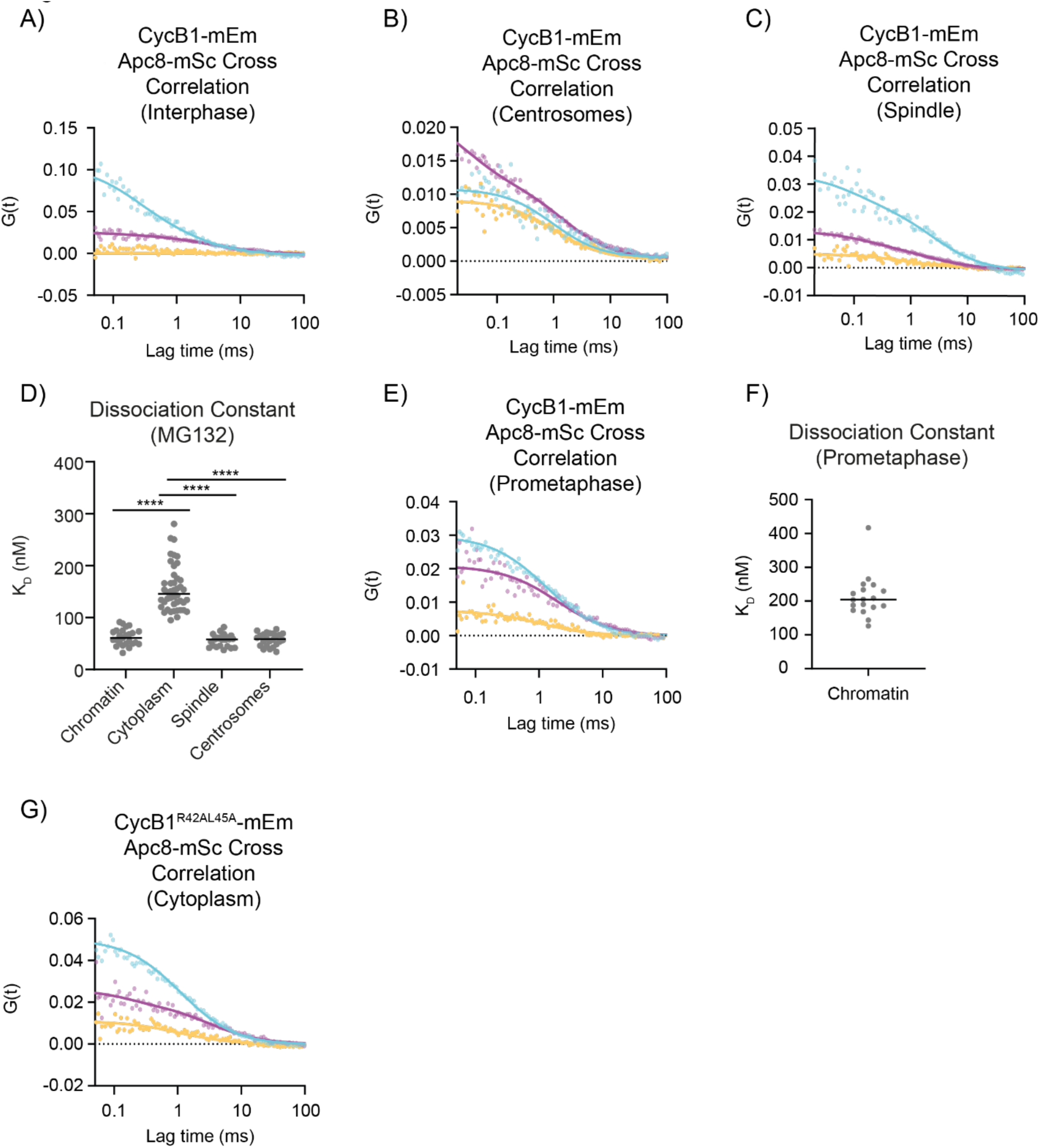
FCCS of Cyclin B1-mEmerald and APC8-mScarlet. A), B), C), E), G) Representative graphs of the autocorrelation function of mEmerald and mScarlet and the cross-correlation function between the two in RPE-1 CCNB1-mEmerald^+/+^; APC8-mScarlet^+/+^ (A,B,E) or in RPE-1 APC8-mScarlet^+/+^ ectopically expressing Cyclin B1^R45AL45A^-mEmerald (G). D), F) Dot plots representing the K_D_ between endogenous Cyclin B1-mEmerald and APC8-mScarlet in metaphase cells following MG132 treatment (D) or in untreated prometaphase cells (F). n ≥ 18 cells per condition, N = 3 independent experiments.

**Figure S3.**
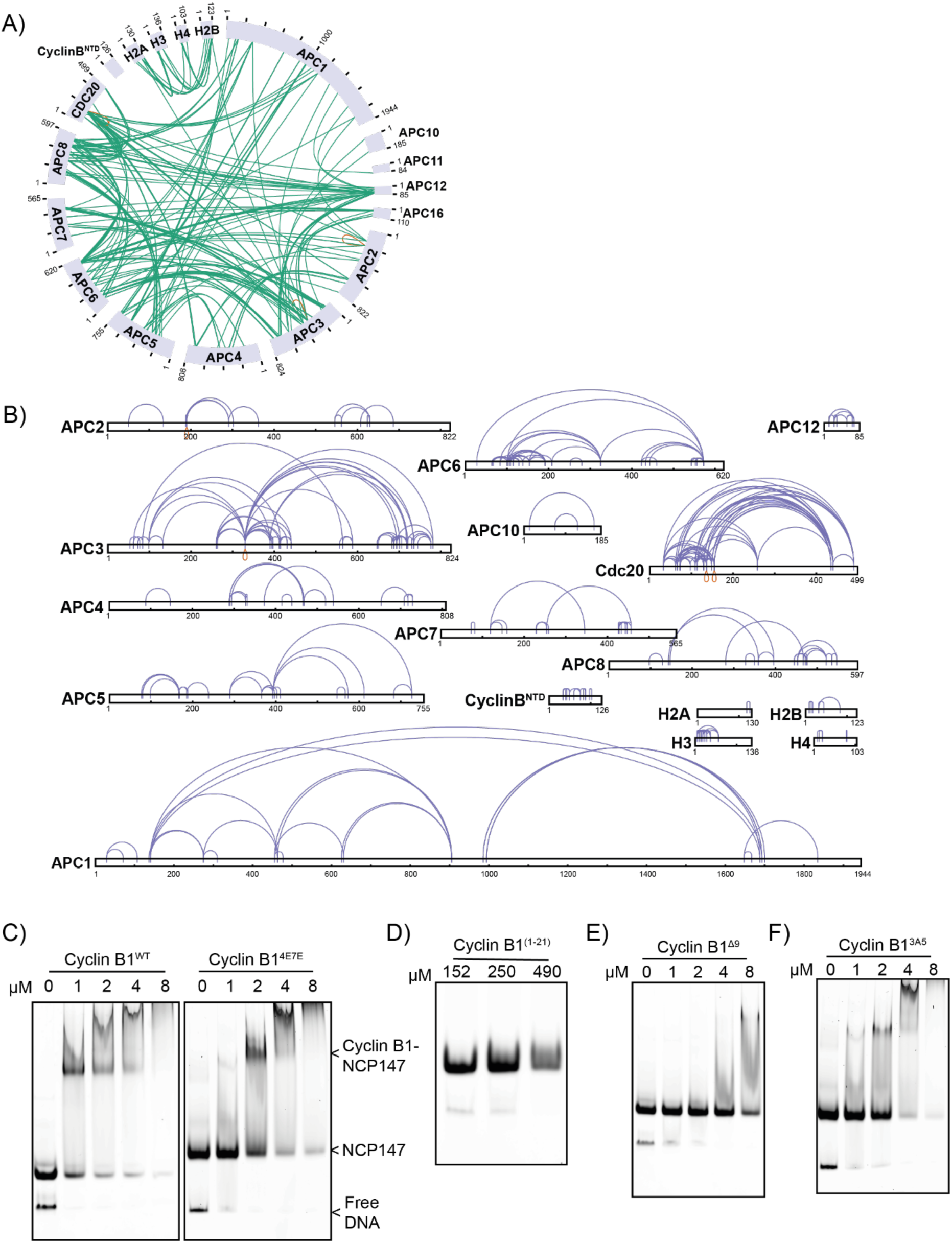
Cyclin B1 binding to nucleosomes. A) Representation of heteromeric crosslinks of the APC/CCDC20-Cyclin B1^NTD^ complex with the NCP. B) Representation of the self-crosslinks of individual APC/C subunits, CDC20, Cyclin B1^NTD^ and the four histones. C,D,E,F). EMSA of NCP 147 with the indicated variant of Cyclin B1. N= 3 independent experiments.

**Figure S4.**
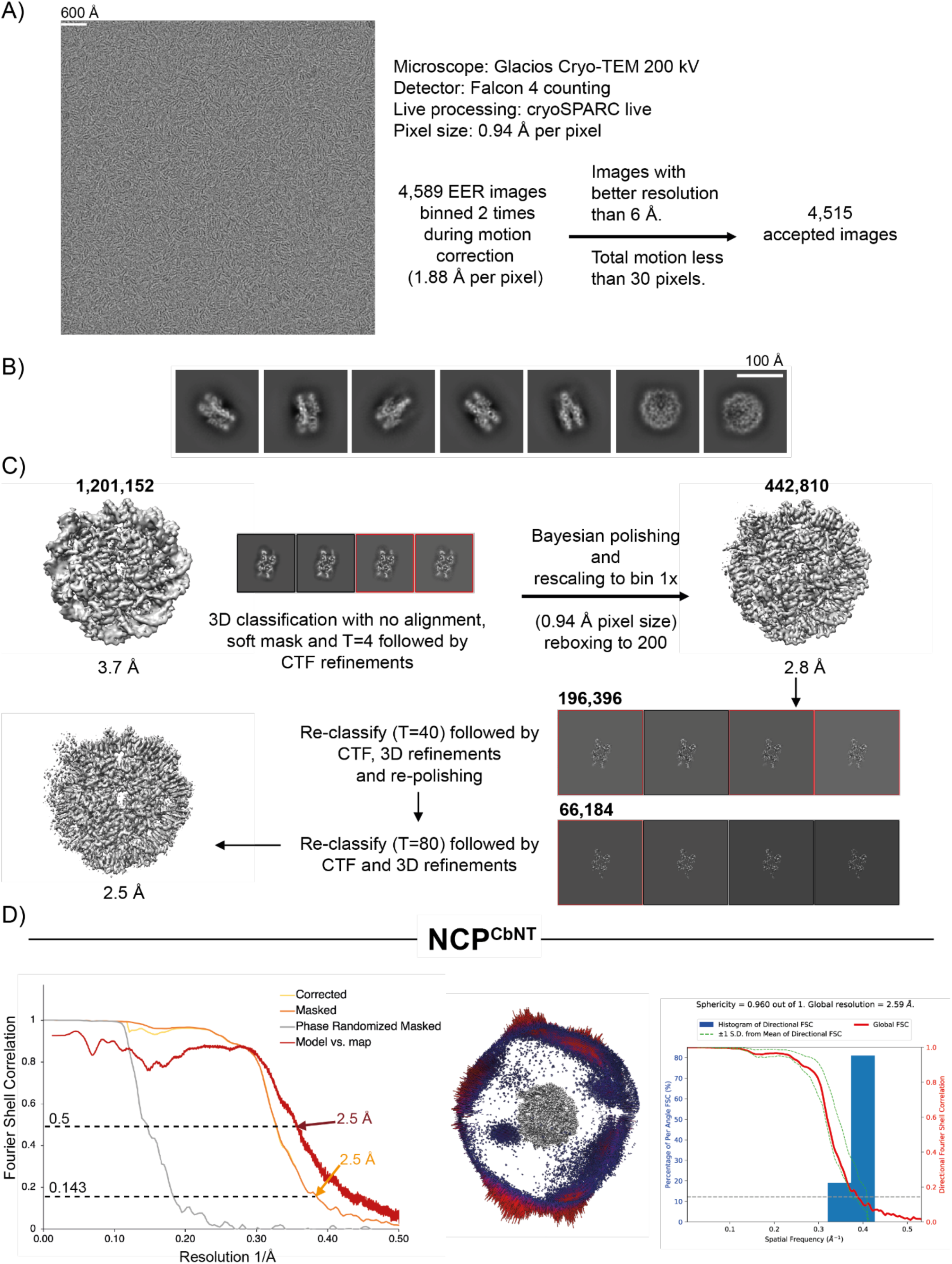
Cryo-EM analysis of the nucleosome core particle in complex with Cyclin B N-terminus. A-C) Workflow showing a representative micrograph, the cryo-EM data collection parameters (A) and the single-particle analysis pipeline for the nucleosome core particle in complex with Cyclin B N-terminus (NCP^CbNT^) (B-C). N. of particles at each classification step is indicated. D) Fourier Shell Correlation (FSC) curves, angular distribution plot and plot of the directional FSC that represents a measure of directional resolution anisotropy for all the reconstructions are shown. Directional FSC and sphericity determination was performed with the 3DFSC software.

**Figure S5.**
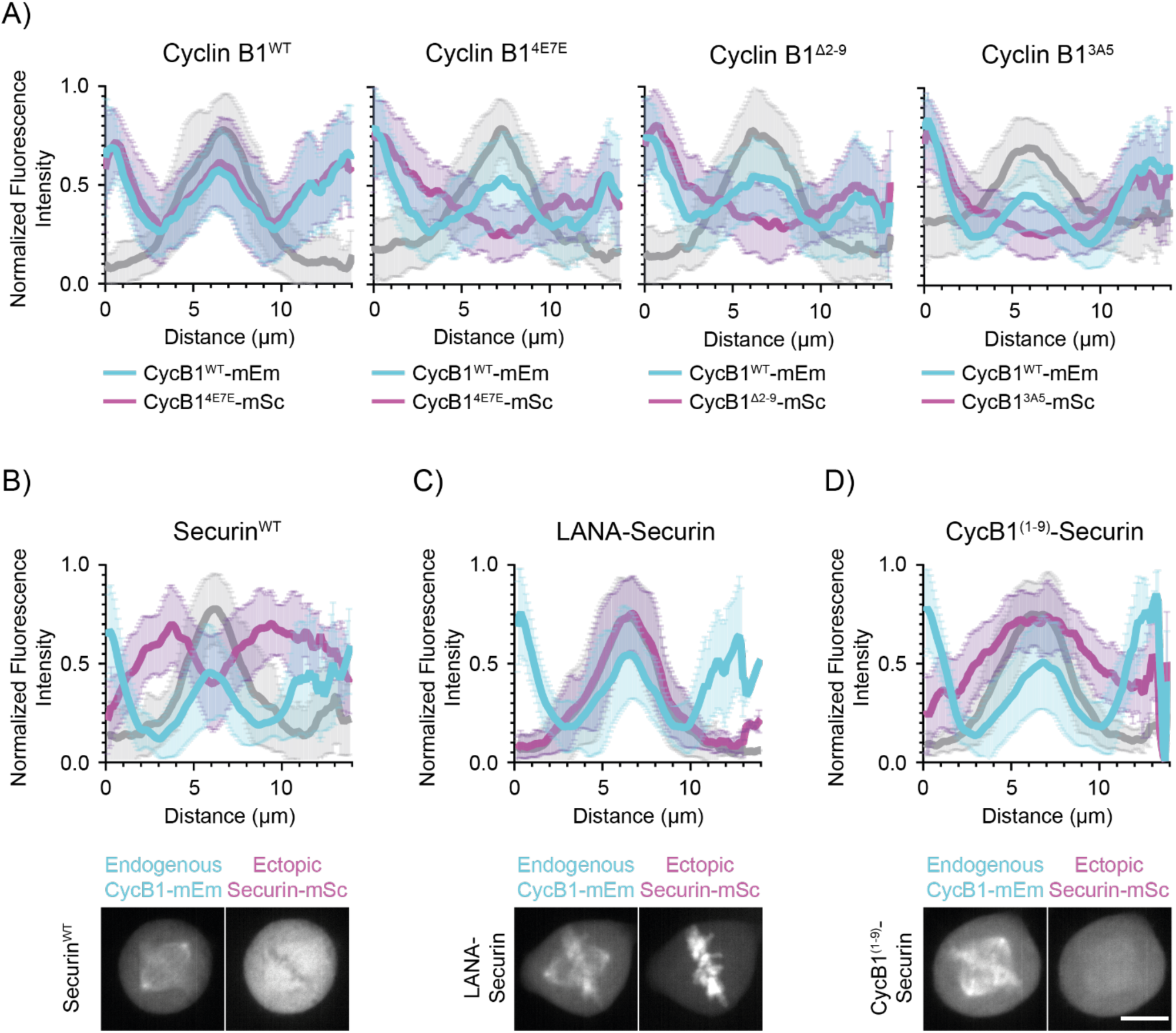
Subcellular localization of Cyclin B1^Δ2-9^ and Cyclin B1^3A5^. A) Graphs representing the pixel-by-pixel fluorescence intensity over a line going from centrosome to centrosome of RPE-1 Cyclin B1-mEmerald^+/+^ cells ectopically expressing the indicated variant of Cyclin B1-mScarlet. Cyan: endogenous Cyclin B1-mEmerand, Magenta: ectopically expressed Cyclin B1-mScarlet, Grey: siR-DNA. N ≥ 21 cells per condition, N = 3 independent experiments. B, C, D) Top: Graphs representing the pixel-by-pixel fluorescence intensity over a line going from centrosome to centrosome of RPE-1 Cyclin B1-mEmerald^+/+^ cells ectopically expressing the indicated variant of Securin-mScarlet. Cyan: endogenous Cyclin B1-mEmerand, Magenta: ectopically expressed Securin-mScarlet, Grey: siR-DNA. Bottom: Maximum projections of representative confocal images used for line profile quantification. Scale bar represents 10 μm. N ≥ 16cells per condition, N = 3 independent experiments.

**Figure S6.**
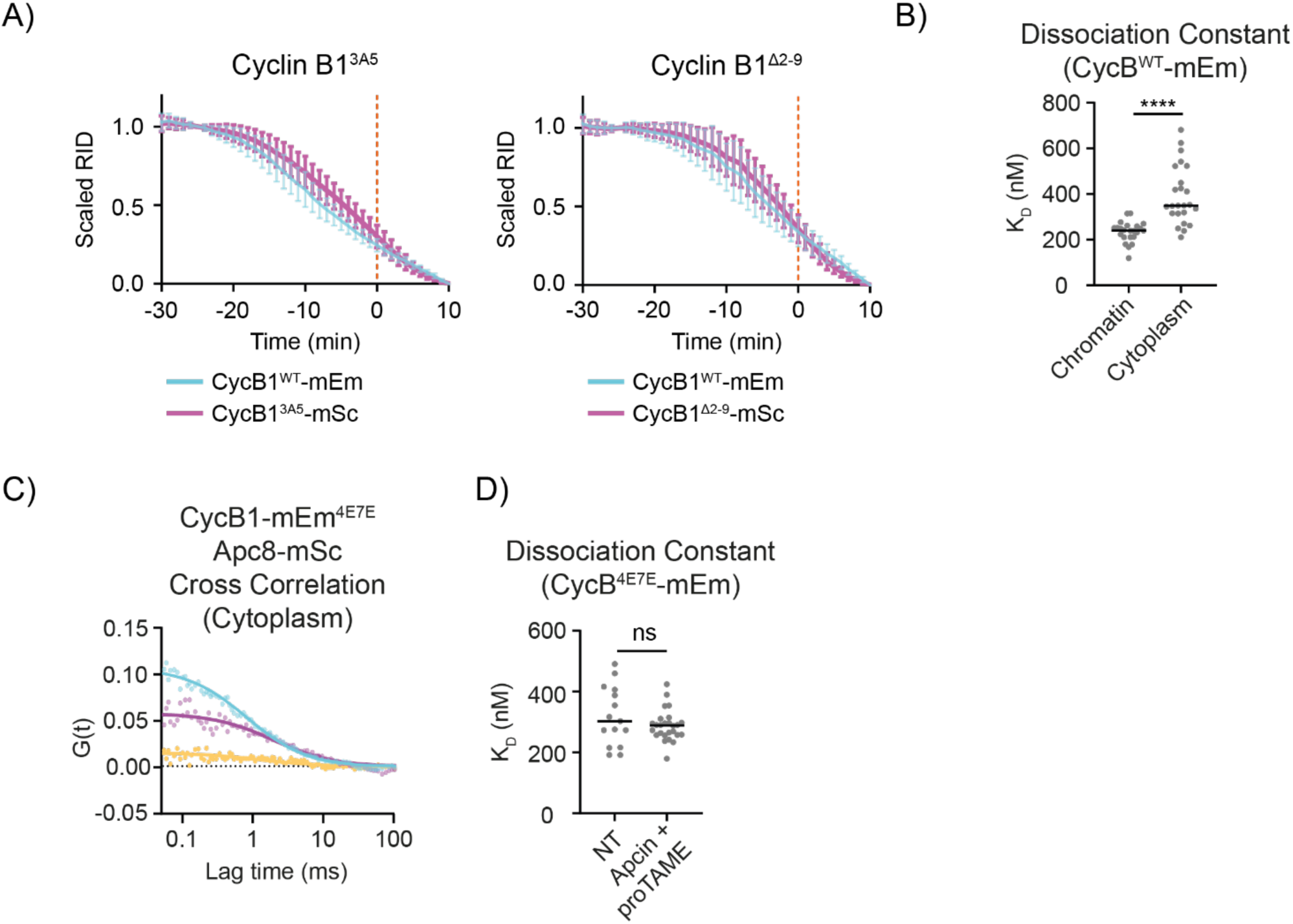
Degradation of Cyclin B1 mutants and APC8 interaction. A) Cyclin B1 degradation graphs representing the fluorescence intensity of Cyclin B1 over time for cells ectopically expressing the indicated Cyclin B1 variants. n ≥ 16 cells per condition, N ≥ 3 independent experiments. B) Dot plot representing the KD values measured for Cyclin B1 ^WT^-mEmerald and APC8-mScarlet by FCCS at different subcellular locations. n = 23 cells, N = 3 independent experiments. C) Representative graph of the autocorrelation function of mEmerald and mScarlet and the cross-correlation function between the two in RPE-1 APC8-mScarlet^+/+^ ectopically expressing Cyclin B1^4E7E^-mEmerald. D) Dot plot representing the KD values measured for Cyclin B1 ^4E7E^-mEmerald and APC8-mScarlet by FCCS before and after a treatment with APCin and proTAME. n ≥ 15 cells, N = 3 independent experiments.

**Figure S7.**
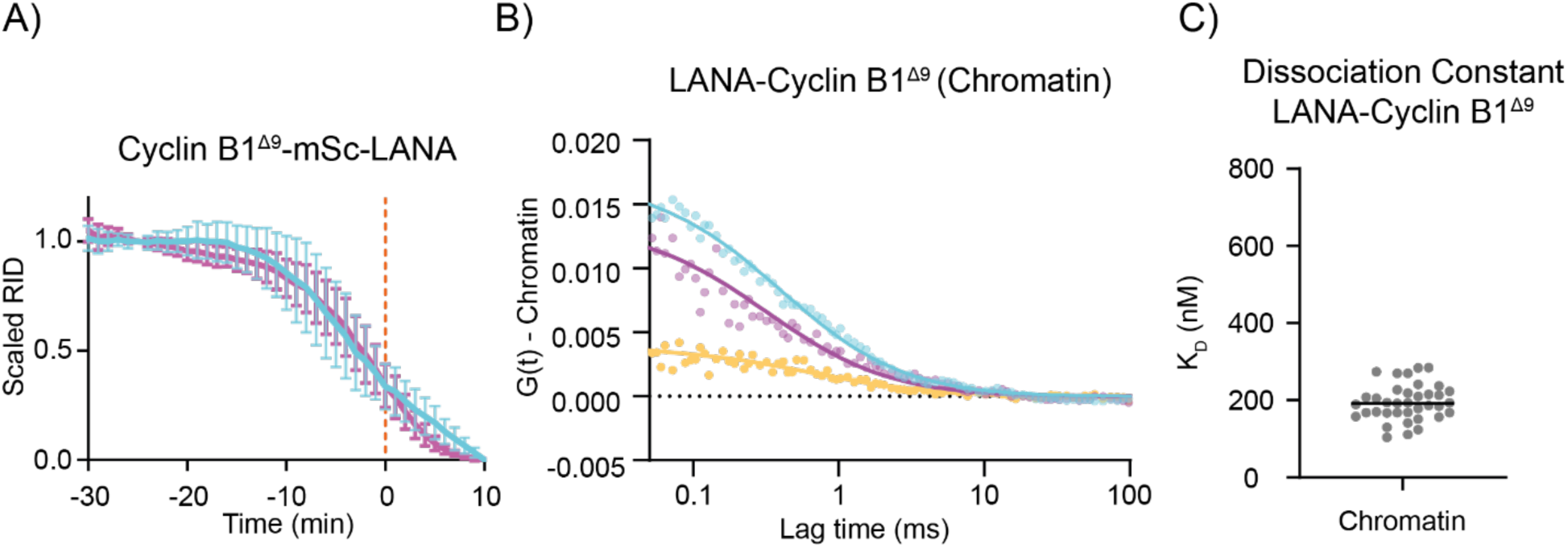
FCCS of LANA-Cyclin B1^Δ9^ -mEmerald and APC8-mScarlet interaction. A) Cyclin B1 degradation graphs representing the fluorescence intensity of Cyclin B1 over time for cells ectopically expressing the indicated Cyclin B1 variants. n = 19 cells per condition, N = 3 independent experiments. B) Representative graph of the autocorrelation function of mEmerald and mScarlet and the cross-correlation function between the two in RPE-1 APC8-mScarlet^+/+^ ectopically expressing LANA-Cyclin B1^Δ9^-mEmerald. C) Dot plot representing the KD values measured for LANA-Cyclin B1^Δ9^-mEmerald and APC8-mScarlet by FCCS in metaphase cells. For all panels, N = 3 independent experiments.

**Figure S8.**
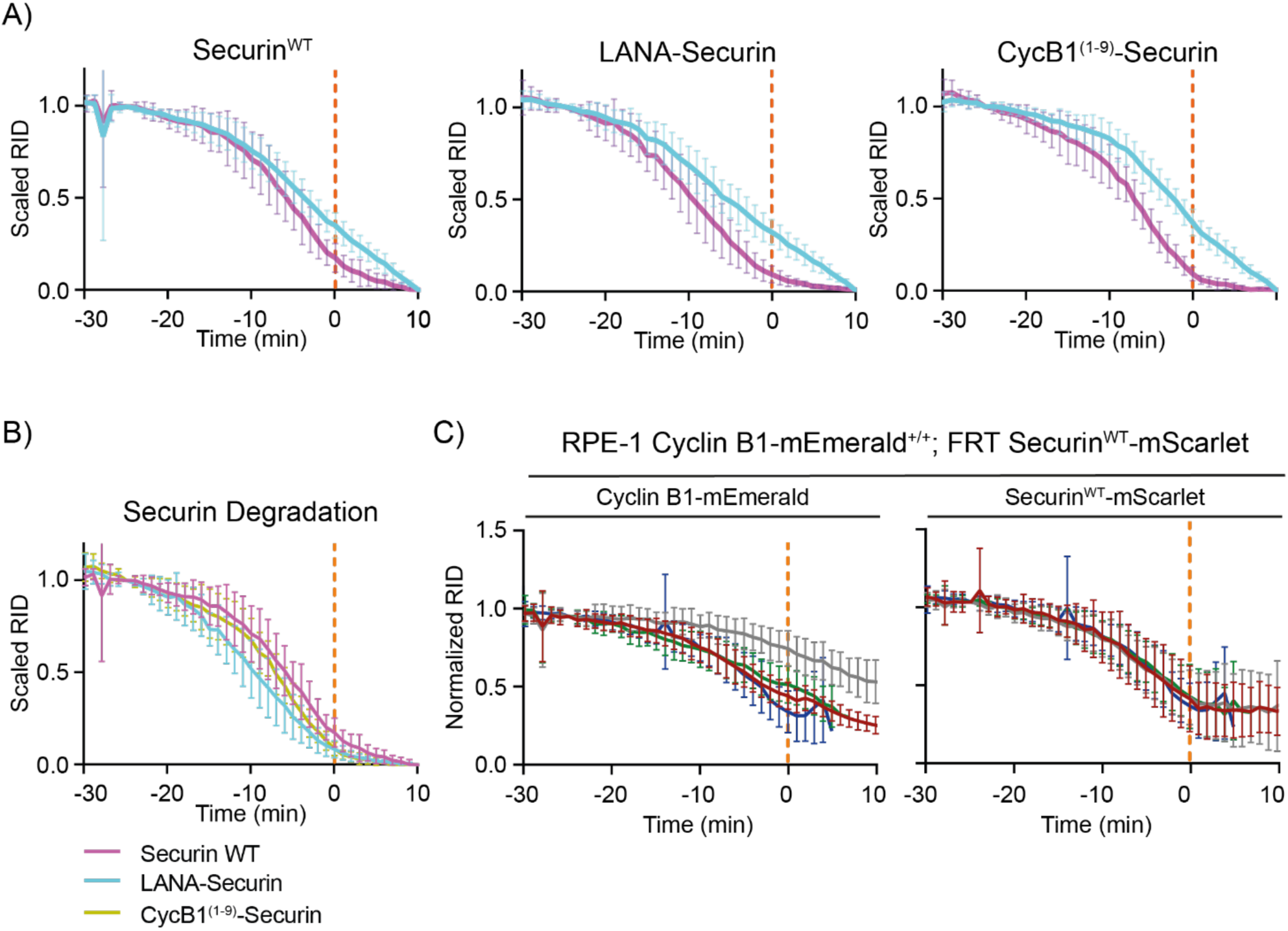
Localizing Securin at the chromatin enhance its degradation. A) Securin degradation graphs representing the fluorescence intensity of Cyclin B1 over time for cells ectopically expressing the indicated Securin variants. Cyan: endogenous Cyclin B1-mEmerald, magenta: ectopic Securin. n ≥ 8 cells per condition, N = 3 independent experiments. B) Securin degradation graph directly comparing the data for Securin degradation shown in A). C) Quantification of normalised Cyclin B1 and Securin fluorescence levels over time measured in RPE-1 CCNB1-mEmerald^+/+^ cells ectopically expressing Securin-mScarlet. N = 12 cells, N = 3 independent experiments.

**Figure S9.**
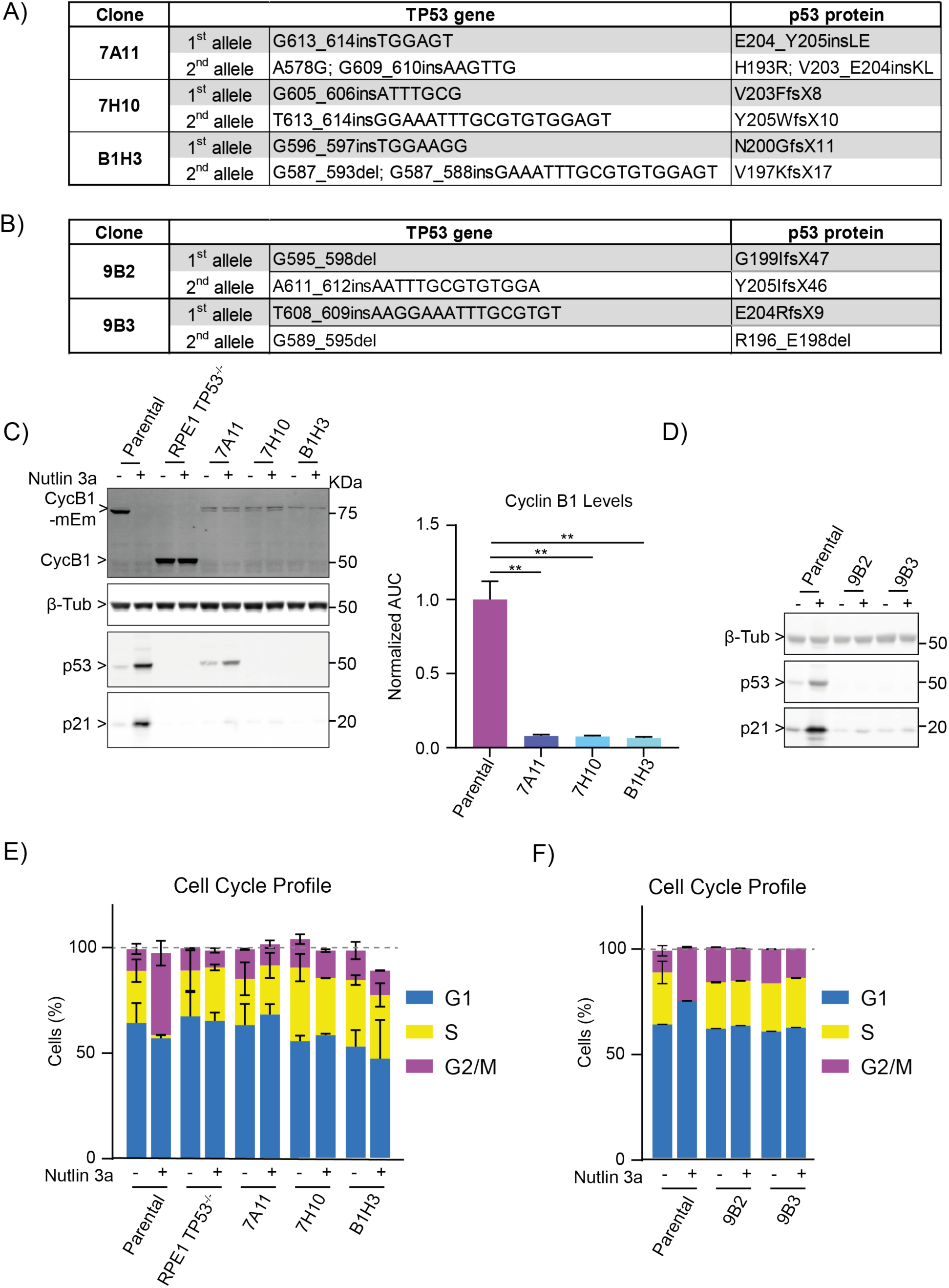
Characterization of RPE-1 CCNB1^4E7E^-mEmerald^+/+^; TP53^-/-^ cells. A) Table recapitulating the sequencing of TP53 gene in RPE-1 CCNB1^4E7E^-mEmerald^+/+^ clones. B) Table recapitulating the sequencing of TP53 gene in RPE-1 CCNB1-mEmerald^+/+^ clones. C) Right: representative immunoblot of cell lysates from parental RPE-1 CCNB1-mEmerald^+/+^, RPE-1 TP53^-/-^ or RPE-1 CCNB1^4E7E^-mEmerald^+/+^ clones, either treated for 24h with Nutlin3a, or left untreated. Left: Bar graph representing the quantification of the immunoblot of Cyclin B1. N = 2 independent experiments. D) Representative immunoblot of cell lysates from parental RPE-1 CCNB1-mEmerald^+/+^ or RPE-1 CCNB1-mEmerald^+/+^; TP53^-/-^ clones, either treated for 24h with Nutlin3a, or left untreated. E) Bar graph representing the quantification of the cell cycle profiles of parental RPE-1 CCNB1-mEmerald^+/+^, RPE-1 TP53^-/-^ or RPE-1 CCNB1^4E7E^-mEmerald^+/+^ clones, either treated for 24h with Nutlin3a, or left untreated. N = 2 independent experiments. F) Bar graph representing the quantification of the cell cycle profiles of parental RPE-1 CCNB1-mEmerald^+/+^ or RPE-1 CCNB1-mEmerald^+/+^; TP53^-/-^ clones, either treated for 24h with Nutlin3a, or left untreated. N = 2 independent experiments. Figure S9 – Supplementary text *Nutlin-3A increased p21 levels in the parental cells, and Cyclin B1 levels decreased as a result of the consequent cell cycle arrest. We did not detect significant changes in p21 or Cyclin B1 in any of the 4E7E p53^-/-^ mutant clones, nor in control RPE-1 TP53^-/-^ cells (Chiang et al., 2019) (Fig. S9C, D). Flow cytometry analysis of PI-stained cells indicated a robust accumulation of 2n and 4n parental cells following a Nutlin-3A treatment, indicative of a p53 dependent cell cycle arrest, whereas RPE-1 TP53^-/-^, Cyclin B1^4E7E^ clones and TP53^-/-^ clones display no signs of cell cycle arrest (Fig. S9E, F). In unperturbed conditions, Cyclin B1^4E7E^ clones and TP53^-/-^ clones did not show any significant variation in cell cycle progression (Fig. S9E, F)*.

**Figure S10.**
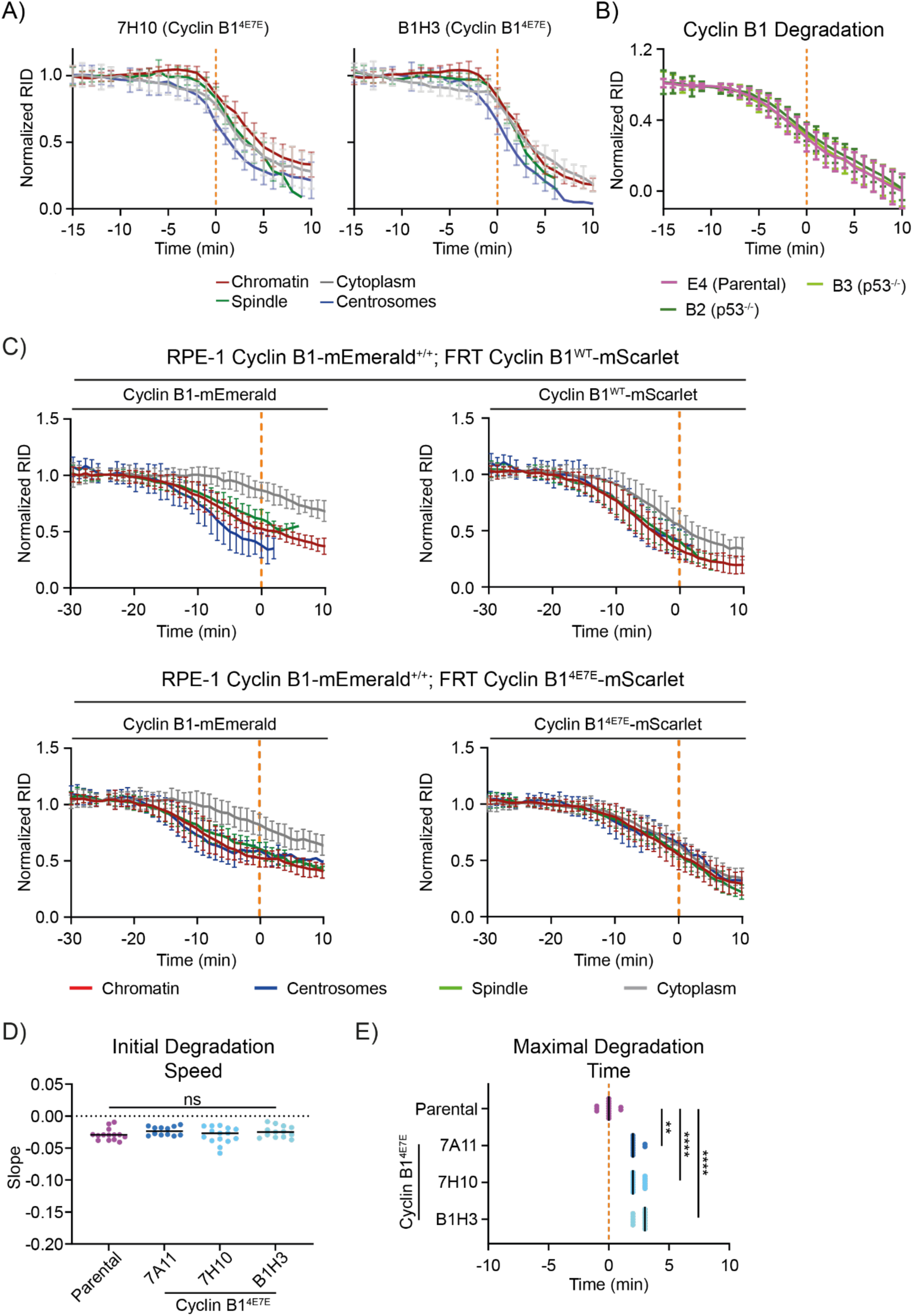
Degradation analysis of Cyclin B1^4E7E^. A, B) Cyclin B1 degradation graph representing the fluorescence intensity of Cyclin B1 over time of RPE-1 Cyclin B1-mEmerald^+/+^ compared to Cyclin B1^4E7E^ clones or p53^-/-^ clones. C) Quantification of normalised Cyclin B1 fluorescence levels over time measured by spinning disk fluorescence microscopy in RPE-1CCNB1-mEmerald^+/+^ cells ectopically expressing the indicated Cyclin-mScarlet B1 variant. n ≥ 18 cells per condition, N = 3 independent experiments. D, E) Dot plots representing the initial degradation speed (D) or the maximal degradation time (E) of RPE-1 Cyclin B1-mEmerald^+/+^ compared to Cyclin B1^4E7E^ clones.

**Table S1.** Concentrations of Cyclin B1-mEmerald and Apc8-mScarlet at different subcellular locations.

| Subcellular Location | Concentration (apparent)<br>of CyclinB1-mEmerald | Concentration (apparent)<br>of APC8-mScarlet |
| --- | --- | --- |
| Chromosome | 120 ± 13 nM | 103 ± 10 nM |
| Cytoplasm | 59 ± 8.5 nM | 99 ± 12 nM |
| Spindle Fibre | 92 ± 7.5 nM | 103 ± 11 nM |
| Spindle poles | 141 ± 17 nM | 98 ± 11 nM |

**Table S2.** Statistical analysis used through the study.

| Table S2 |  |  |  |  |  |
| --- | --- | --- | --- | --- | --- |
| Figure | Mean difference<br>(p value) |  | Multiple comparison<br>(p value) |  | Statistical Test |
| Fig 1H | <0.0001 |  | Cytoplasm vs. Chromatin | <0.0001 | Kruskal-Wallis; Dunn's Multiple Comparison |
|  |  |  | Cytoplasm vs. Spindle | <0.0001 |  |
|  |  |  | Cytoplasm vs. Centrosomes | <0.0001 |  |
| Fig 1I | 0.3435 |  |  |  | one-way ANOVA |
| Fig 1J | 0.2348 |  |  |  | unpaired t-test |
| Fig 7E | <0.0001 |  | Parental vs 7A11 | <0.0001 | Welch's ANOVA; Dunn's Multiple Comparison |
|  |  |  | Parental vs 7H10 | <0.0001 |  |
|  |  |  | Parental vs B1H3 | <0.0001 |  |
| Fig 7F | 0.075 |  |  |  | Kruskal-Wallis |
| Fig 7G | <0.0001 |  | Parental vs 7A11 | 0.016 | Kruskal-Wallis; Dunn's Multiple Comparison |
|  |  |  | Parental vs 7H10 | <0.0001 |  |
|  |  |  | Parental vs B1H3 | <0.0001 |  |
| Fig S1C | top left | 0.936 |  |  | one-way ANOVA |
|  | top right | 0.629 |  |  | one-way ANOVA |
|  | bottom left | 0.304 |  |  | one-way ANOVA |
|  | bottom right | 0.464 |  |  | one-way ANOVA |
| Fig S1D | top | 0.797 |  |  | two-way ANOVA |
|  | bottom | 0.9201 |  |  | Kruskal-Wallis |
| Fig S1F | left | 0.379 |  |  | one-way ANOVA |
|  | middle | <0.001 |  |  | one-way ANOVA |
|  | right | 0.2 |  |  | Kruskal-Wallis |
| Fig S2D | <0.0001 |  | Cytoplasm vs. Chromatin | <0.0001 | Kruskal-Wallis; Dunn's Multiple Comparison |
|  |  |  | Cytoplasm vs. Spindle | <0.0001 |  |
|  |  |  | Cytoplasm vs. Centrosomes | <0.0001 |  |
| Fig S6B | <0.0001 |  |  |  | unpaired t-test |
| Fig S6D | 0.1952 |  |  |  | unpaired t-test |
| Fig S9C | 0.001 |  | Parental vs. 7A11 | 0.0016 | one-way ANOVA; Tukey's Multiple Comparison |
|  |  |  | Parental vs. 7H10 | 0.0016 |  |
|  |  |  | Parental vs. B1H3 | 0.0016 |  |
| Fig S10D | 0.234 |  |  |  | Kruskal-Wallis |
| Fig S10E | <0.0001 |  | Parental vs 7A11 | 0.01 | Kruskal-Wallis; Dunn's Multiple Comparison |
|  |  |  | Parental vs 7H10 | <0.0001 |  |
|  |  |  | Parental vs B1H3 | <0.0001 |  |

**Table S3.** Cryo-EM data collection, refinement and validation statistics.

| Table S3 |  |
| --- | --- |
|  | NCP-CbNT |
| Data collection and processing |  |
| Magnification | 150.000 |
| Voltage (kV) | 200 |
| Electron exposure (e-/Å<sup>2</sup>) | 60 |
| Defocus range (µm) | -0.6 to -1.6 |
| Pixel size (Å) | 0,94 |
| Symmetry imposed | C1 |
| Initial particle images (no.) | 1.201.152 |
| Final particle images (no.) | 38.352 |
| Map resolution (Å) | 2,5 |
| FSC threshold | 0,143 |
| Refinement |  |
| Initial model used (PDB code) | 3LZ0 |
| Model resolution (Å) | 2,5 |
| Map sharpening B factor (Å<sup>2</sup>) | -22 |
| Model composition |  |
| Nonhydrogen atoms | 11895 |
| Protein residues | 748 |
| Nucleotides | 290 |
| Ligands | MN(8), CL(2) |
| B factors (Å<sup>2</sup>) |  |
| Protein (min/max/mean) | 11.50/79.20/37.16 |
| Nucleotide (min/max/mean) | 17.98/201.32/83.66 |
| Ligand (min/max/mean) | 30.66/106.65/46.81 |
| R.m.s. deviations |  |
| Bond lengths (Å) | 0,005 |
| Bond angles (°) | 0,704 |
| Validation |  |
| MolProbity score | 2,07 |
| Clashscore | 9,36 |
| Poor rotamers (%) | 4,99 |
| Ramachandran plot |  |
| Favored (%) | 97,8 |
| Allowed (%) | 2,2 |
| Disallowed (%) | 0 |

## Notes

### Competing Interest Statement

The authors have declared no competing interest.

